# The formation of an expanding memory representation in the hippocampus

**DOI:** 10.1101/2023.02.01.526663

**Authors:** Sachin P. Vaidya, Guanchun Li, Raymond A. Chitwood, Yiding Li, Jeffrey C. Magee

## Abstract

How brain networks connected by labile synapses store new information without catastrophically overwriting previous memories remains poorly understood^1,2^. To examine this, we tracked the same population of hippocampal CA1 place cells (PC) as mice learned a task for 7 days. We found evidence of memory formation as both the number of PCs maintaining a stable place field (PF) and the stability of individual PCs progressively increased across the week until most of the representation was composed of long-term stable PCs. The stable PCs disproportionately represented task-related learned information, were retrieved earlier within a behavioral session, and showed a strong correlation with behavioral performance. Both the initial formation of PCs and their retrieval on subsequent days was accompanied by prominent signs of behavioral timescale synaptic plasticity (BTSP), suggesting that even stable PCs were re-formed by synaptic plasticity each session. Further experimental evidence supported by a cascade-type state model indicates that CA1 PCs increase their stability each day they are active eventually forming a highly stable population. The results suggest that CA1 memory is implemented by an increase in the likelihood of new neuron-specific synaptic plasticity, as opposed to extensive long-term synaptic weight stabilization.

## Introduction

Beneficial information learned by animals during a behavioral episode should be retained and later retrieved during similar experience to improve behavior^3,4^. In the hippocampus, a brain area thought to be involved in episodic memory^5-9^, learning during an experience produces environment-specific neural activity (context discriminability)^10-13^ that includes high densities of PCs at reward sites and salient feature locations (over-representations) ^9,14-18^. How this information is retained across time remains an open question as PC coding across days has been reported to be noisy with minor, if any, statistical structure, or conserved elements^14,19-28^. Neuronal tagging studies (e.g., c-Fos) provide some evidence that memory retrieval, and by implication, storage may occur within active hippocampal PCs^5-7^ but the principles of stable memory formation remain elusive.

The generation of experience-specific hippocampal PC activity patterns during learning is currently thought to involve alterations to the synaptic weights within the main subregions (CA3 & CA1) via a plasticity form known as behavioral timescale synaptic plasticity (BTSP) ^29-34^. The long-term stabilization of these learned weights could underlie memory formation. However, a complexity that arises with extensive synapse stabilization is that although it facilitates long-term information storage by protecting against subsequent overwriting, it can quickly saturate network capacity and limit additional learning in neural networks^35,36^. Thus, some balance of synaptic stability and plasticity must be achieved within networks to facilitate continual learning^1,2,37^. One proposed solution for this stability-plasticity dilemma is for synapses to have multiple stability states that they can transition through in an experience-dependent manner, with only a small fraction of synapses ultimately reaching high levels of stability (sparseness) ^38-40^. Currently, it remains unknown how neuronal networks in general and the hippocampus in particular maintain stable memory storage in the face of continuous learning.

### Neural representations reflect task-related information

To examine this issue, we used two-photon Ca^2+^ imaging to longitudinally record the activity of a single hippocampal CA1 neuronal population for 7 days as head-fixed transgenic mice expressing GCaMP6f^41^ learned two separate reward locations on a linear treadmill that was enriched in tactile features (2511 total pyramidal neurons tracked; 126±5 laps/session; 70 sessions over 7 days in 10 mice) (Fig. 1a; Extended Data Fig. 1).

**Fig. 1:**
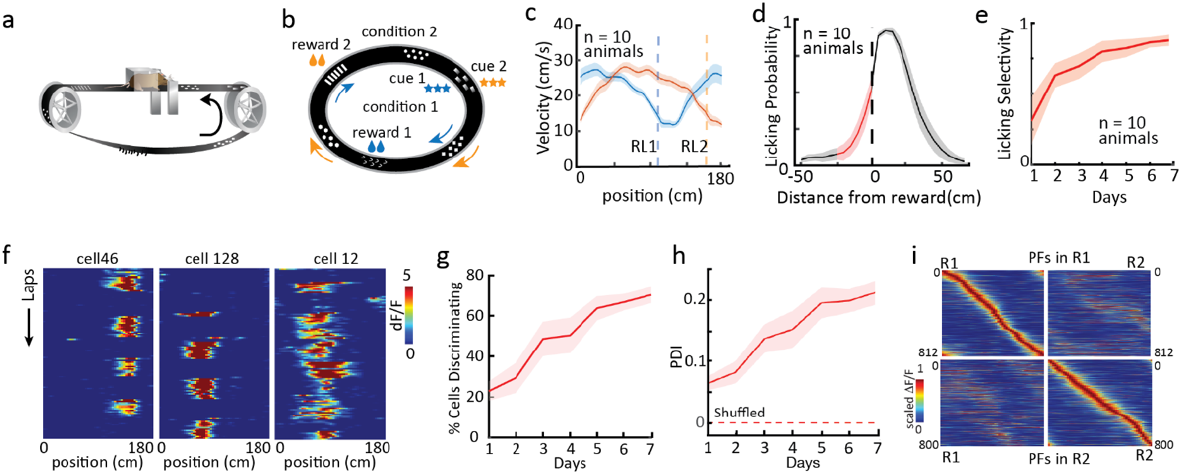
Daily evolution of behavior, single neuron and population activity. **a**, Schematic of experimental apparatus and (**b)** two different cue-reward location conditions. **c**, Average velocity profiles for mice during condition RL1 (blue; shading is SEM) and RL2 (orange) on day7. Note selective slowing near reward locations. **d**, Licking probability in reference to reward locations with anticipatory licking shown in red for day 7. **e**, Licking selectivity (see Methods) as a function of behavior day. **f**, example PCs with RL2 preference, RL1 preference, and no preference (left to right). **g**, Percentage of PCs showing high cell discriminability increased across experimental days. **h**, Population discriminability index (PDI) increases with experimental day. Dashed line shows PDI for shuffled data. **i**, Heat maps of PCs with PF activity in condition RL1 (top), during RL1 (top left) and RL2 (top right; both sorted by activity during RL1). Bottom, same as top but for PC with PFs active in condition RL2 (sorted by activity during RL2).

The mice were first habituated on a featureless track as a sucrose-water reward was delivered at random locations. On day 1, the habituated mice were exposed to a new feature-containing track with two alternating reward locations given in 12-18 lap blocks. Reward location was contingent on a specific light cue given to either the left or right eye at a set location (Fig. 1b). Behavioral performance, as assessed by running and licking patterns improved over the 7 days (Fig. 1c-e). Both single neuron (Fig. 1f&g; Extended Data Fig. 2a) and population (Fig. 1h; Extended Data Fig. 2b-d) PC activity also evolved over the recording days such that by day 7 two highly distinct and stable neuronal activity patterns had developed (discriminability), each with its own PC spatial density profile (over-representations) (Fig. 1i; Extended Data Fig. 2e-g). In addition, we observed a subtle increase in the day-to-day correlations of the PC representation across the week that potentially hints at a stabilization process occurring during repeated learning (Extended Data Fig.2h-j). These data are similar to that found in most previous studies that have used comparable recording techniques and behavioral conditions^14,20,22,24,27, but see 28^.

### Emergence of a stable memory representation with experience

To look for the formation of a memory representation in this neuronal population we identified active PCs for each reward location (RL) on each experimental day for the entire week (see methods for PC criteria). The total population of active PCs for any given day was relatively constant (average PC count/day from 10 mice combined=767±8 PCs, n=14; for 2 RL conditions over 7 days; PCs are 30.5% of imaged pyramidal neurons). CA1 neurons showed a varying degree of PC activity across days, with most cells having no PFs and a smaller fraction having PF activity across multiple days. This long-tail distribution of PF activity was markedly different from what would be expected if all PCs had an equal chance of PF formation each day (Fig.2a,b)^21^. A closer look revealed that the history of prior PC activity was a strong determinant of PF formation on subsequent days. Cells with established PFs on prior days were not only more likely to form new PFs but tended to do so at the same location whenever they reappeared (Fig.2c-d).

We next systematically followed the activity of PCs across the week and found that the percentage of PCs active on Day 1 that maintained a consistent PF location for consecutive days decreased progressively from ∼35% on the next day (day 2) to ∼6% by day 7 (PF on each subsequent day within ±30 cm of location on the first day; see methods and Extended Data Fig. 3). Performing this same analysis for each of the subsequent days (i.e., days 2-7) showed a similar rate of decrease in PC counts for each day (Fig. 2e). The decay of consistently active PCs that first appeared on days 1&2 were well fit by double exponential functions (t_fast_=0.67±0.02 day; t_slow_ =4.6±0.19 days; n=4, 2 RL conditions for each day; Fig. 2e; Extended Data Fig. 3). The differential decay in PC populations was reflective of enhanced stability in a subset of PCs^19-24^, suggesting that PCs could be distinguished based on the duration of their sustained activity across days. The presence of PCs with enhanced stability (presumably those producing τ_slow_) was further evident in the accumulation of PCs that were previously present (past PCs), while the number of newly appearing PCs (new PCs) decreased through the week (Fig. 2f-h). The quantity of past PCs was higher than that expected from a random memoryless process (data; past PC total=2284 vs expected=490; Fig. 2h; Extended Data Fig. 3).

**Fig. 2:**
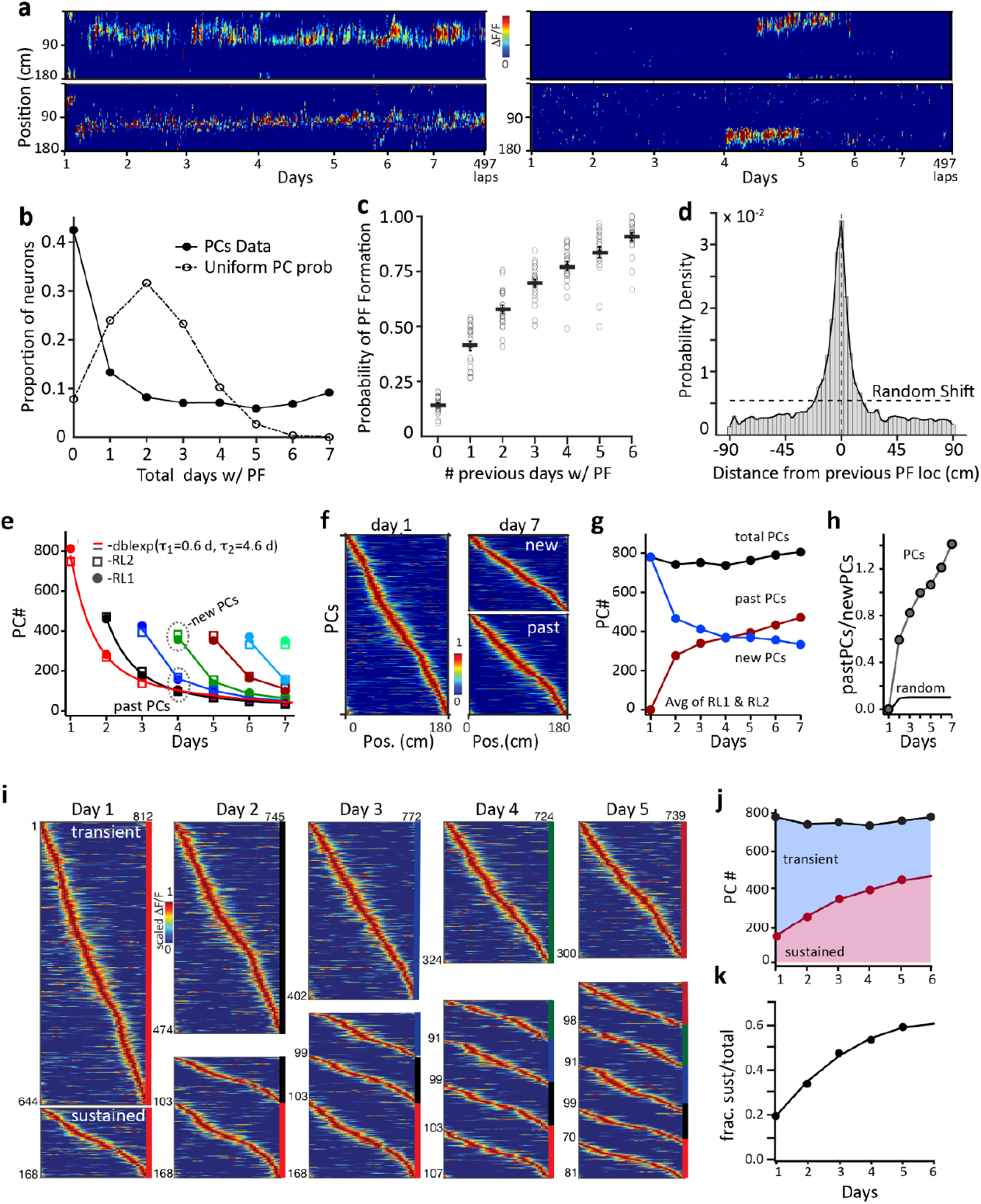
Emergence of a stable memory representation. **a**, Example cells with varying PC activity across days. **b**, Distribution of PC activity across 7 days; solid: Actual PC data with long-tail distribution (average RL1 and RL2); dotted: Expected distribution with uniform PC formation probability each day. **c**, Probability of PC formation as a function of the number of days the PC was previously active; data from all 7 days sorted together. (n=20, 10 animals, RL1 and RL2 separate) **d**, The distance from the previous PF appearance if a PC reappears on a subsequent day (n=7853 reappearances (RL1 and RL2 combined). **e**, counts for PCs that first appear on day 1 and maintained a consistent PF on subsequent days (consistent PF present on each day from 1 through 7; red). Other colors are PCs that first appear on days 2-7 and maintained a consistent PF on subsequent days (day 2, black; day 3, dark blue; day 4, dark green; day 5, dark red; day 6 light blue; day 7, light green). PCs during RL1 (squares) and RL2 (circles). Solid lines over PC counts for days 1 and 2 are double exponential functions produced by fits of data. **f**, (left) PCs active on day 1 sorted by PF location, (right) PCs active on day 7 separated into new PCs (upper; have not maintained consistent PFs) and past PCs (lower; maintained consistent PFs from previous days), sorted by PF location on day 7. **g**, PCs divided into past PCs (red) and new PCs (blue) plotted against experimental day. All PCs (which is equal to the sum of past and new PCs; black) versus experimental day. PC counts are the average of RL conditions. **h**, Ratio of past PC to new PC counts for each experimental day (Gray circles). Calculated ratio from random process (black line; Extended Data Fig. 3). **i**, Visualization of transient PCs and sustained PCs evolution for each experimental day using PC activity heat maps for each group sorted by PF location in that population. Color bars on sides are coded according to (e). **j**, PC counts for sustained group (red circles) and total PC count (black circles). Solid line (red) is a projection using a double exponential fit. Blue shading is transient model pool count and red shading is sustained pool count. **k**, Fraction of total PCs count that are sustained PCs versus experimental days (circles). Solid line is a projection using a double exponential fit.

The two different decay rates described above suggest a relatively straightforward method for isolating PCs into separate populations based on the number of days that they maintained consistent PFs (transient PCs ≤ 2 days; sustained PCs > 2 days). Separation of PCs using this criterion revealed the accumulation of PCs within the sustained group at the expense of the transient population (Fig. 2i-k). Indeed, the proportion of the total active PC population provided by sustained PCs increased nearly three-fold after 5 days of repeated learning and this was significantly correlated with a similar increase in the stability of the total PC population over the same days (Fig. 2j; Extended Data Fig. 4a-c). Together, these data suggest that PC formation on each day involves a plastic group that rapidly becomes inactive or has an unstable PF tuning, and a more stable population that forms the basis of a progressively expanding memory representation of the animal’s past experience over time.

### Sustained Place Cells encode learned task information

For the sustained PCs to function as a memory of what was previously learned we would expect to find properties of the sustained representations that reflect the animals prior experience. Thus, we determined the spatial density profiles of the two groups of PCs (transient PCs < 2 days; sustained PCs > 2 days to minimize classification error; Fig.3a) and found that the PC density (PC count/cm) around salient regions (reward and cue) was substantially more elevated in the sustained group than in the transient population (see confidence intervals in Fig. 3b, Extended Data Fig. 4d,e). In addition, we determined the level at which each individual neuron (Cellular Discrimination Index; CDI) as well as the entire population (Population Discrimination Index; PDI) was able to discriminate the two different reward conditions and found that sustained PCs were more discriminative than transient (mean CDI; sustained:0.172±0.003, n=2455 cells, transient:0.089±0.002 n=1874 cells, p=<1.0e^-7^ unpaired two-tailed students t-test; mean PDI; sustained: 0.14±0.02, n=10 mice, transient: 0.09±0.01 n=10 mice, p=1.0e^-3^ paired two-tailed students t-test; Fig. 3c-e, Extended Data Fig. 4f). Thus, the experience-dependent development of both the spatial over-representation of salient regions and reward condition discrimination were found to be markedly elevated in the sustained over the transient PC group.

**Fig. 3:**
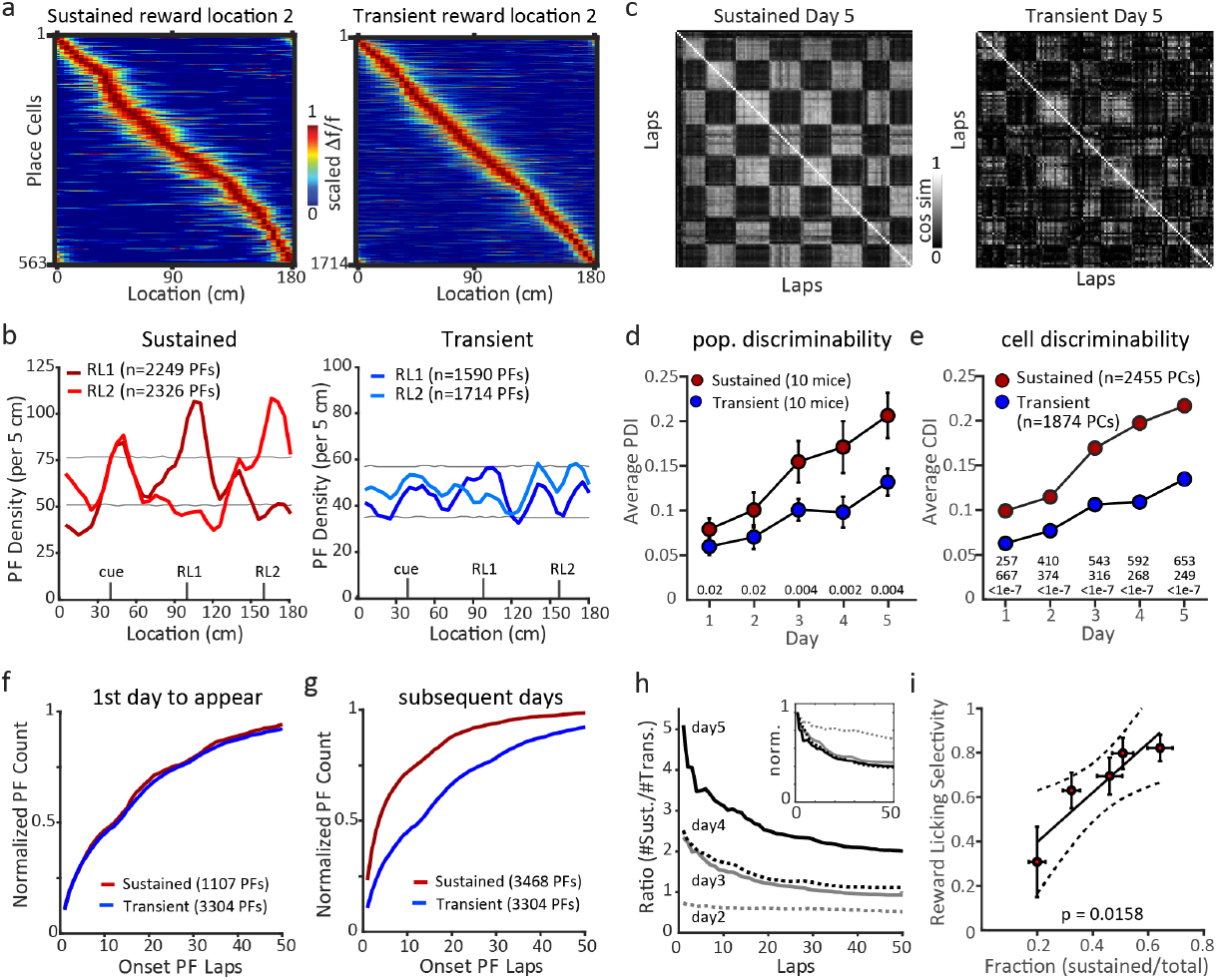
Properties of sustained and transient PC groups. **a**, PC activity heatmaps for sustained (left) and transient (right) groups under RL2 condition days 1-5 combined. **b**, PF densities for sustained PCs (left, red) and transient PCs (right, blue) under RL1(dark colors) and RL2 (light colors) conditions. Gray lines are 99% confidence intervals of random distribution produced via bootstrapping 10,000 times. Sustained groups include identified PFs from days 1-7 and transient groups includes identified PFs from days 1-5. **c**, Cosine similarity matrices for sustained (left) and transient (right) populations on day 5 from a single mouse. **d**, Population discriminability (PDI) versus experimental day, p-values from two-way, paired t-tests are shown for each day. **e**, cell discriminability (CDI) versus experimental day, n- (sustained, upper; transient, middle) and p-values (lower) from two-way, unpaired t-tests are shown for each day. **f**, Cumulative distribution of onset lap (first trial PC appeared) for the first days that PC appeared for sustained (days 1-5) and transient groups (days 1-6). **g**, Sustained group on subsequent days (2-7). Transient group is same as in panel (**f). h**, Ratio of number of sustained PCs to number of transient PCs for a given lap for days 2-5 (gray dashed day 2; gray solid, day 3; black dashed, day 4; black solid, day 5). Inset shows ratios normalized to the peak amplitude of each day. **i**, Reward licking selectivity versus fraction of total population that is composed of sustained PCs. P-value for linear regression is shown.

We next asked if the sustained population was active over the same range of trials during a given behavioral session as the transient population. We found that while the range of PC onset trial was similar between the two groups on the first day that a PC appeared (KS test p = 0.9541; Fig. 3f, Extended Data Fig. 4g), on all subsequent days the sustained PCs became active much earlier in the session than the transient population (KS test p = 3.63e-06; Fig. 3g; Extended Data Fig. 4h). This caused the fraction of the total representation contributed by the sustained population to be higher at the beginning of each session, particularly on later days (Fig. 3h). Thus, stable PCs appear to be immediately retrieved into activity during each new day’s experience. Finally, we found that the daily increase in the selectivity of licking that occurred across the week of behavior was highly correlated with the proportion of the total PC population made up of sustained PCs on a given day (Fig. 3i, Extended Data Fig. 4i). These data suggest that the daily rapid retrieval of a growing memory representation could contribute to the progressive enhancement of the behavior observed in these mice.

### Place Cells are established and daily reconstituted by BTSP

Accumulating evidence suggests that behavioral timescale synaptic plasticity (BTSP) is the primary mechanism of PC formation and learning-related changes in CA1 population activity^29-34^. BTSP is a directed form of synaptic weight plasticity that is induced when input from the entorhinal cortex (EC3) drives Ca^2+^ plateau potentials in the dendrites of CA1 neurons. Thus, we next examined the activity of all PCs for the known signatures of BTSP^29-32^, which include an abrupt PF appearance that was associated with a large amplitude Ca^2+^ signal, as expected for the strong burst firing driven by dendritic plateau potentials, and a backward shifted PF, whose width correlated with the velocity at plateau induction. (Fig. 4a, e-g). The fraction of neurons with detected BTSP events as well as the number of observed plateaus per cell was significantly higher in sustained PCs than transient PCs on the first day of appearance. (Fig. 4b; 5-day mean, sustained; 5.40 ±0.27 plateaus/session; transient 3.55±0.17 plateaus/session, (n = 100 sessions); p=<1.0e^-15^, paired two-tailed students t-test; see Extended Data Fig.5 for calibration and confirmation studies for plateau detection criteria). We also observed that the rate of putative plateau potentials increased with the number of days a given neuron exhibited PC activity (Extended Data Fig 6), which also correlated well with the observed increase in the probability of PF formation (see Fig. 2C; R^2^=0.76, p∼0.00, data from RL1 and RL2, 10 animals).

**Fig. 4:**
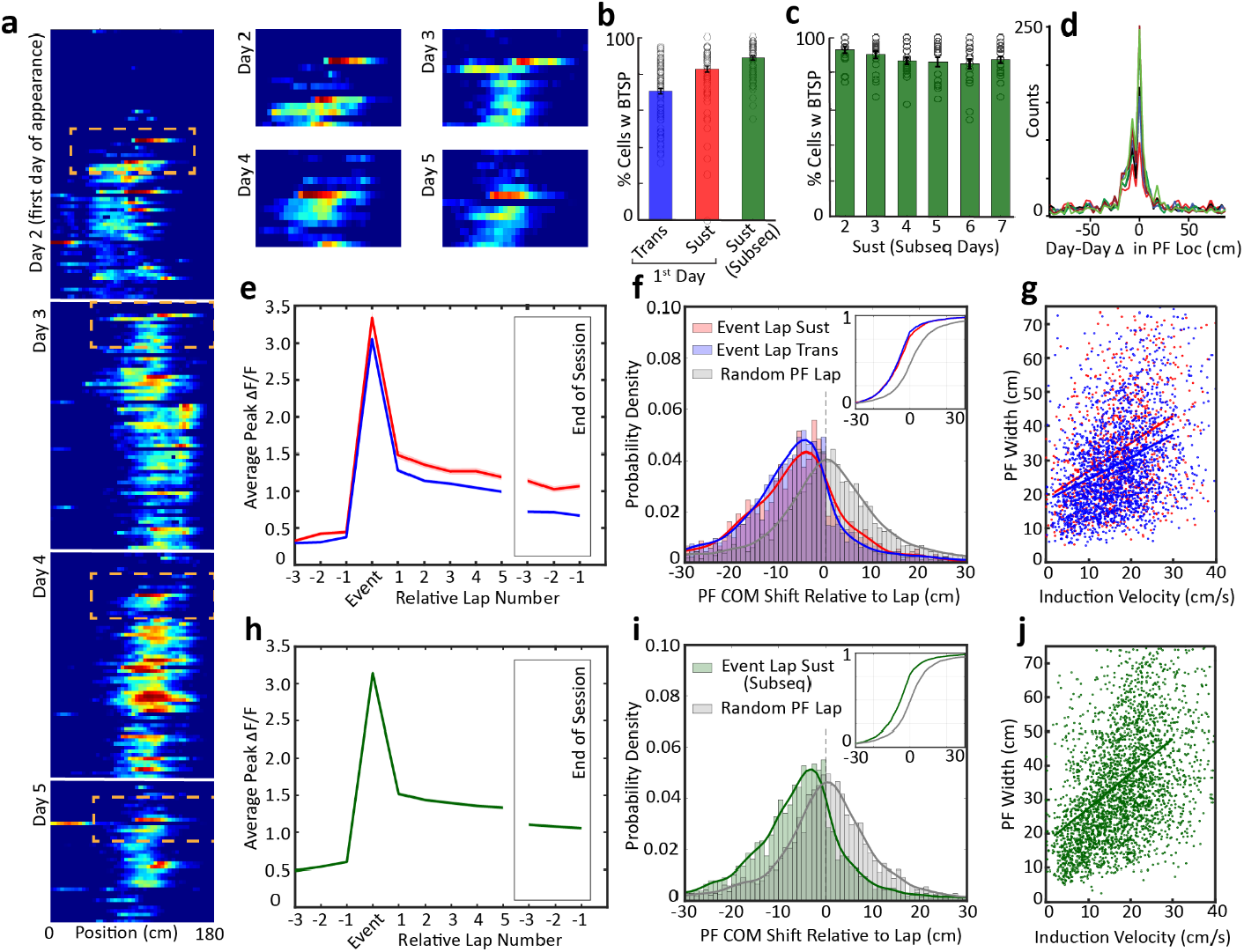
Place Fields are established and daily reconstituted by BTSP. **a**, Example of a sustained cell from its first day of appearance, day 2 through day 5 with first detected putative plateau potentials inset. **b**, Percentage of cells per session detected with BTSP (see methods) on first day of transient (blue) and sustained (red) cell appearance and when sustained cells were subsequently active. Data from 100 sessions (10 animals, Day 1-5, RL1 and RL2; 80 sessions, Day 2-5 for equivalent sessions for Subsequent). **c**, Percentage of cells per session with detected BTSP for sustained cells that were reconstituted for each day (20 sessions per day, 10 animals, RL1 and RL2). **d**, Predictive skew in distribution of day-to-day PF location shifts, RL1 and RL2 combined. **e-g**, Signatures of BTSP in transient (blue, n= 2324 events) and sustained (red, n = 934 events) PFs on first day of appearance with (e) abrupt establishment of a PF after a strong Calcium event, (f) a negative predictive PF shift in COM for laps following the Calcium event (gray denotes a random PF firing lap for comparison, Sustained p = 4.65e-17 KS test, Transient p = 5.57e-42, KS test, p=0.19 Random Sustained vs Random Transient KS test) and (g) correlation between velocity in the onset lap and PF width (Sustained : y = 0.80 x + 18.95, p = 1.13e-32; Transient y = 0.64 x + 18.23, p = 2.1e-62). **h-j** same as e-g but for reconstitution of sustained PFs (n = 3045 events) on subsequent days of activity (Shift: Event lap vs Random lap p = 8.1e-62, KS test; Ind Vel vs PF Width: 1.02 x + 17.27, p = 9.61e-155).

Remarkably, the signatures of BTSP induction were also prominent in almost all of the sustained PCs on each of the subsequent days that they were active (Fig. 4b,c) and this was so common that a pronounced backward shift in the day-to-day location of sustained PCs was observed at the population level (Fig 4d). Moreover, this plateau induced synaptic plasticity appeared to be needed for robust PF firing on each subsequent day as only weak residual PC activity, if any at all, was observed at the start of the session (fig 4.h-j; Extended Data Fig 6). These data suggest that the ‘stable’ PC activity of sustained neurons across days is produced by additional BTSP induction at the same PF location^42^. The somewhat unexpected finding that the stability of sustained PC firing across days is not solely produced on the first day of induction but also requires additional plasticity at the same location suggests that stable memory formation and retrieval is achieved through the daily reconstitution of PCs.

### Cascade model with history dependence captures PC dynamics

To better understand the above PC dynamics, we generated several theoretical models and assessed their ability to adequately capture the properties of observed data. We first examined a competitive interaction between three pools of model “neurons”, which simulated (1) available pyramidal cells, (2) transient cells and (3) sustained cells (see Methods; Extended Data Fig. 7a-e). The different decay rates in the model (based off the double exponential fits in Fig. 2e) produced a sustained group that decayed back to the available population nearly an order of magnitude more slowly (τ_sust_=4.6 days) than the transient population (τ_trans_ = 0.67 days) (Extended Data Fig. 7f&g). The resulting accumulation of the sustained model cell group over 7 days simulated the activity of real PCs and accurately predicted the PC counts over the week (MSE= 182, n=28; Extended Data Fig. 7 h-j). However, an extension of the 3-pool model using individual model neurons could not account for various properties of individual PCs such as the distribution of the number of days a given neuron had PF activity or the probability of reappearance after a missed day of PF activity (Fig.5a, b), and this is most likely due to the lack of any history dependence in PF formation in this model.

Cascade models of synapses featuring a gradient of stability or metaplastic states are known to optimize memory retention in neural networks^38-40^. We followed a similar approach by modeling the formation of PCs across days using a cascade of states that increased the likelihood of PF reconstitution with each day of PF activity. Briefly, following activity on a given day, PCs transitioned either to a state from which their activity and PF location was reliably reconstituted the next day (right side; red arrows, Fig 5c) or to a less reliable state from which they were probabilistically recruited (left side; blue arrows, Fig 5c). The likelihood of an active cell transitioning to the reliable state as well as the probability of being recruited to be active from the unreliable state increased with the number of days a PC was active (Methods, Fig 5c, Extended Data Fig.6). This resulted in a gradient of probability for PF formation based on prior PF history that was similar to that of observed plateau potential rates in the experimental data (Extended Data Fig.6). Moreover, the cascade model adequately captured the influence of history not only with the distribution of days with PF activity but also in the probability of reappearance for previously active PCs when they missed a single day of activity (Fig. 5a,b). The latter result along with the observation that experimental data could be reproduced by a model featuring no backward transitions suggest that the decay times for the observed stability are substantially longer than one week. The probabilistic reconstitution of PFs in the cascade model replicated the observed slow decay of PCs maintaining the same location of PF activity over days as well as the gradual buildup of sustained PCs over time (Fig 5d,e). An alternative model with progressive stability where decay rates decrease with each day a PC is active accounted for PF counts across days but failed to replicate the distribution of days with PF activity (Extended Data Fig.8). These results suggest that during repeated learning hippocampal CA1 representations are progressively stabilizing both at the population and individual neuron levels and that this stabilization is based on past experience. Moreover, evidence suggests that the mechanism of this PC stabilization may be an enhanced probability of dendritic plateau potential initiation and subsequent BTSP induced PC reconstitution.

**Fig. 5:**
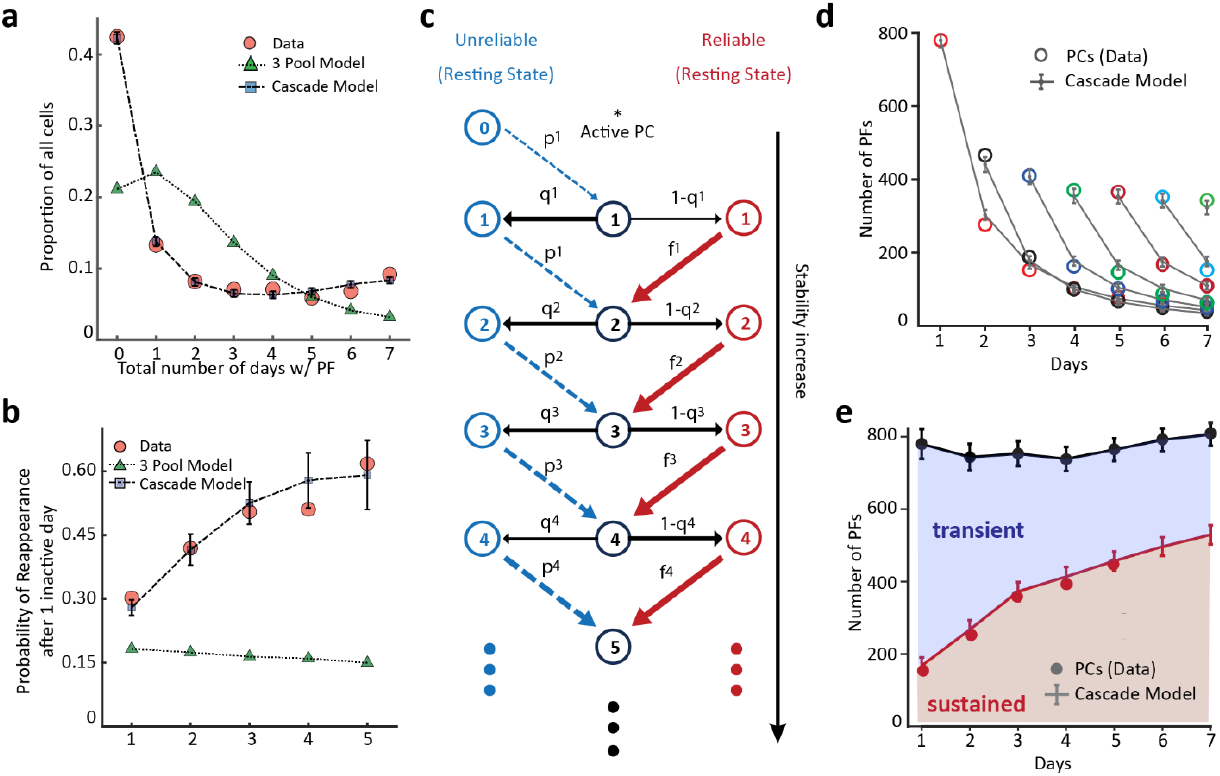
A Cascade model with history dependence captures PC dynamics. **a**, Distribution of the proportion of cells that had PF activity for a given number of days, for the experimental data (average RL1 and RL2) and that observed in the 3-pool model and cascade-like state model. **b**, Probability of PF formation for PCs that were inactive for one day but were subsequently active, sorted by days of previous PF activity for the experimental data, 3-pool model and Cascade-like state model. **c**, Schematic of the cascade model. Left (blue) and Right (red) side signify the resting state of the PC in between days with PF activity. Centre (black) depicts days with active PF activity. **d&e**, Cascade model adequately captures the distribution of sustained and transient cells observed in experimental data across days.

## Discussion

### Summary

The evidence presented here indicates that CA1 representations are stabilized over the course of repeated learning as the likelihood that particular PCs will exhibit consistent PF activity increases continuously. The long-term stable population of PCs encodes learned task information better than the less stable population as stable PCs are biased to the salient regions of the environment and have higher context discriminability. This stable population is also retrieved earliest in each behavioral session and a strong correlation between the size of the stable group and behavioral performance suggests that the immediate access to learned information from the animal’s past is useful for performance. Interestingly, we found evidence that these stable cells are reconstituted every day by BTSP, just as in de novo PC formation. Finally, a cascade-type state model that accounts for past PC activity by progressively increasing the probability of subsequent PF formation adequately explains our experimental findings adding further support to the idea that PC stability increases with each day of activity to yield a population of highly stable PCs. Together these data indicate that the size of a highly stable, information rich, readily retrievable PC representation that is behaviorally useful grows with repeated learning and that the cellular and circuit level mechanisms may be somewhat unconventional given the current theory that memory formation is achieved via an extensive, long-term synaptic weight stabilization^46,47^. Instead we find evidence that substantial weight decay occurs overnight with subsequent new plasticity required for robust PF activity which is conceptually similar to mechanisms used in memory models proposed to deal with the large amount of instability inherent within nervous systems^48-50^.

### Comparison with past results

Numerous labs have tracked CA1 hippocampal PC activity over days and most of them report a progressive decrease in the similarity of population activity from a given day onward (decreases in population vector correlations or increases in decoding error). In general, most PCs only transiently join the representation, but there is usually a second population that maintains consistent PF activity for many days^9,14,19-24,27,43^. This is very similar to what we describe as different pools of PCs that can be distinguished by their stability. Several studies have observed varying degrees of pairwise session-to-session activity correlation changes with level of experience which could indicate changes in the stability of the PC population activity (most show increases^14,20,22,24, 27^; but see^28^). The variability among these studies may be due to differences in recording methods, mouse behavior or level of learning attained by the time of the experiment. A similar analysis of our data produced correlation values and daily changes in them that were within the range of most of these studies, suggesting comparable levels of stable/unstable PFs for our conditions (See Extended Data Fig.2). Thus, with some exceptions, we would expect the progressive accumulation of long-term stable PCs that we report here, to also be present in the data of these other groups, even though they have not commented on this aspect.

Two groups report that the average stability of PFs is elevated around reward sites but it is unclear if this is due to an elevated density of stable PCs encoding the salient region or to simply an elevation of the stability of all PCs at that region^9,22^. Interestingly these groups do separate out PCs either into a group experiencing more sharp-wave/ripple (SPWR) activity^22^ or Fos expression^9^ levels and both of these PC groups seem to preferentially encode the non-salient regions of the environment. Finally, the hippocampal population map is known to be generated each day with new PCs being added progressively during a session^30,44^ but we are unaware of any other studies that have compared the rate of PC appearance between stable and unstable PCs. The fraction of PCs exhibiting BTSP signatures during this process that we report here is comparable to that of another study (∼55% versus ∼75%)^29^, and the plateau potential rates we observed are very similar to those previously reported for novel conditions (∼0.07 versus ∼0.08 PP/lap) ^45^. In the end, we think that most of the findings reported in this manuscript will be present within the data collected by other groups with any substantial discrepancies being due to differences in learning behavior or in the PC grouping (SWR activity or Fos expression).

### Increased likelihood of new plasticity versus extensive synaptic weight stabilization

As mentioned above, PF formation and the associated representation learning in the hippocampus is thought to be implemented by BTSP modification of synaptic weights within the main subregions (recurrent CA3 synapses and Schaffer collateral synapses from CA3 to CA1)^29-34^. The classical hypothesis would be that stabilization of learned Schaffer collateral synaptic weights on CA1 PCs produces the long-term storage needed for memory formation and retrieval in this network^46,47^. More recent theoretical work on network models of memory storage have found that it is advantageous to stabilize only a fraction of learned synaptic weights while leaving the majority available for overwriting during continuous learning conditions^36-40^. Indeed, the above evidence suggests that in all PCs, including those maintaining stable representations across days, the majority of synaptic weights modified during learning do not maintain their strength between sessions. This may also include the CA3 network, although previous work suggests that activity there is relatively stable on subsequent days^23^. Either way a large amount of neuron-specific Schaffer collateral weight adjustment appears to be needed to recreate past activity patterns, suggesting that most weights in the network remain fully available for new plasticity with only a sparse set being highly stabilized.

An increased probability of burst firing has been observed in putative memory encoding CA1 neurons^8^ and the correlation between dendritic plateau potential rate and PC stability observed here suggests that the main locus of the long-term stability could be in the likelihood of dendritic plateau initiation. Similarly, given that plateau initiation and the burst firing that it produces, is controlled by various circuit components, including excitation, inhibition and neuromodulation, many elements may be involved here. One possibility is that long-term information storage is achieved not through the stabilization of an extensive fraction of synaptic inputs sufficient to directly drive activity but instead in the conjunction between a small subset of stabilized weights from different input pathways to control the induction of new plasticity (e.g. Schaffer collateral and perforant path synapses). This new plasticity then, since it is both neuron and location specific, reforms past activity patterns during retrieval. It may in fact turn out that the most important stabilization is within the perforant path input, which is a potent regulator of dendritic plateau potentials in CA1 neurons. This is particularly intriguing given previous experimental work on the importance of distal dendrite synapses for learning in the neocortex and theoretical work on the capabilities of weight adjustments to synaptic inputs from teaching signals that target distal dendrites^51-54^. Finally, neuron-specific dendritic inhibition and modulation mechanisms may also be involved^55-57^ and future work examining these theoretical and experimental ideas is needed.

In sum, our observations suggest that the increasing stabilization of certain PCs produces an informative, readily retrievable, and progressively expanding memory representation in the mouse hippocampus that when retrieved is useful for behavior. Here, memory formation may proceed through the progressive stabilization of a sparse set of synapses that instead of directly driving activity, induces new plasticity that then recovers past activity patterns during retrieval. This may represent an efficient mechanism based on past experience for the encoding and maintenance of episodic memories in the hippocampus^35,58^.

## Acknowledgements

We thank B. Ujfalussy for suggesting progressively stabilizing model schemes, S. Romani for helpful discussions and C. Grienberger, S. Romani and H. Zoghbi for comments on the manuscript. This work was supported by the Howard Hughes Medical Institute and the Cullen Foundation. The data that supports the findings of this study are available from the corresponding author upon request. The code that supports the findings of this study is available from the corresponding author upon request.

## Author contributions

S.P.V., R.A.C. and JCM designed experiments. S.P.V. performed the imaging experiments and S.P.V. and JCM analyzed the data. JCM created the 3-pool model. GL created the cascade model and contributed to the analysis of history dependence. R.A.C. performed slice experiments and analyzed data. Y.L. performed in vivo voltage recordings and Y.L. & JCM analyzed data. S.P.V and JCM wrote the manuscript with comments from all authors.

**Extended Data Fig. 1:**
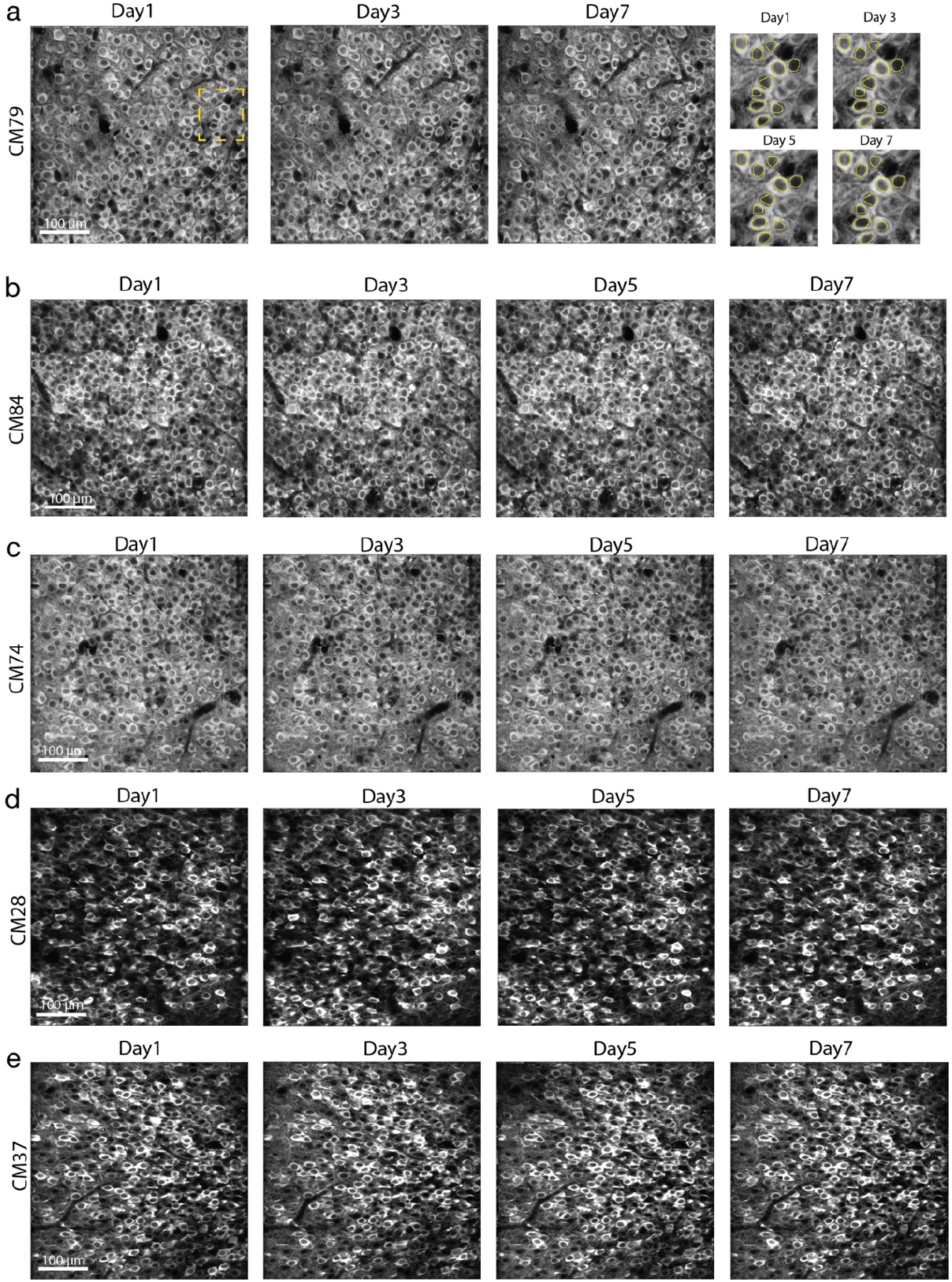
Multi-day imaging of CA1 neuron population. **a-e**, 2-photon imaging Field of View (FOV) of the CA1 area from 5 representative animals across the duration of longitudinal imaging (7 days). The last panel in (**a**) shows representative ROIs used for signal extraction. The representative animals were randomly chosen and each FOV is a z-stack of 500 frames randomly chosen from the middle of the imaging session on each representative imaging day.

**Extended Data Fig. 2:**
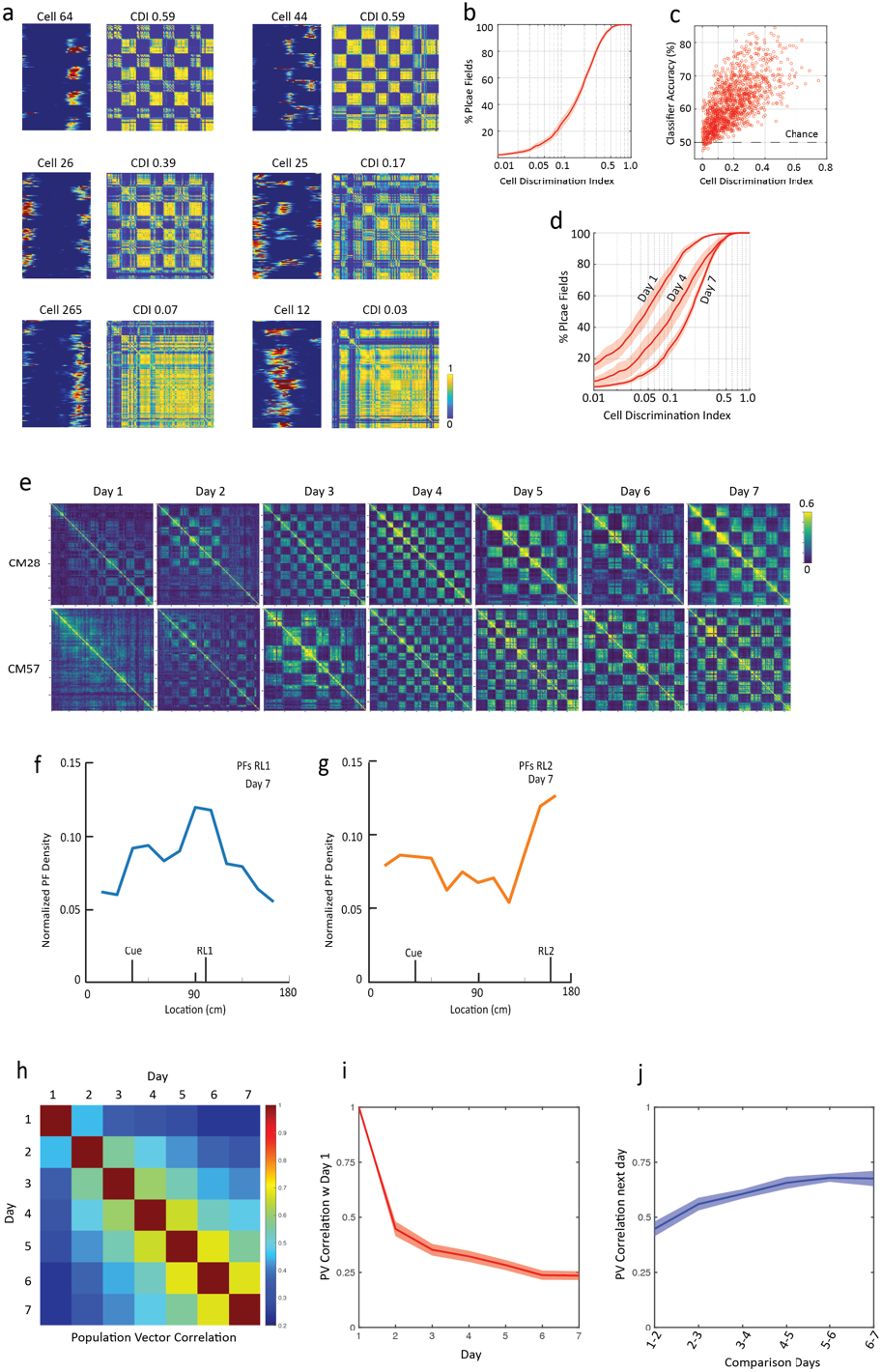
Characterization of single neuron and population activity. **a**, Cosine Similarity Matrices for 6 representative cells with their Cell Discrimination Index (CDI) describing their discriminative ability between RL1 and RL2 (see Methods). **b**, Distribution of CDI values in a given animal on day 7 (n=10 animals). **c**, A discriminability threshold for CDI was established using a supervised learning model on real and shuffled data. Model accuracy shown here for non-shuffled data. The model could discriminate above chance level when CDI of the cell was >0.1. This was set as discriminatory threshold in Fig.1g. **d**, Distribution of CDI values in a given animal across day 1, 4 and 7. **e**, Representative Cosine Similarity Matrices for population data for two animals across 7 days. PDI (Population Discrimination Index) was calculated from these matrices (see Methods). **f & g**, Normalized PF spatial density on day 7 for PFs in RL1 and RL2. Data is from all active PFs. **h**, Population Vector Correlation Matrix calculated as the average of 10 animals (RL1 and RL2) separate (n=20). **i**, PV correlation with activity on day 1. **j**, PV correlation for activity on successive days.

**Extended Data Fig. 3.**
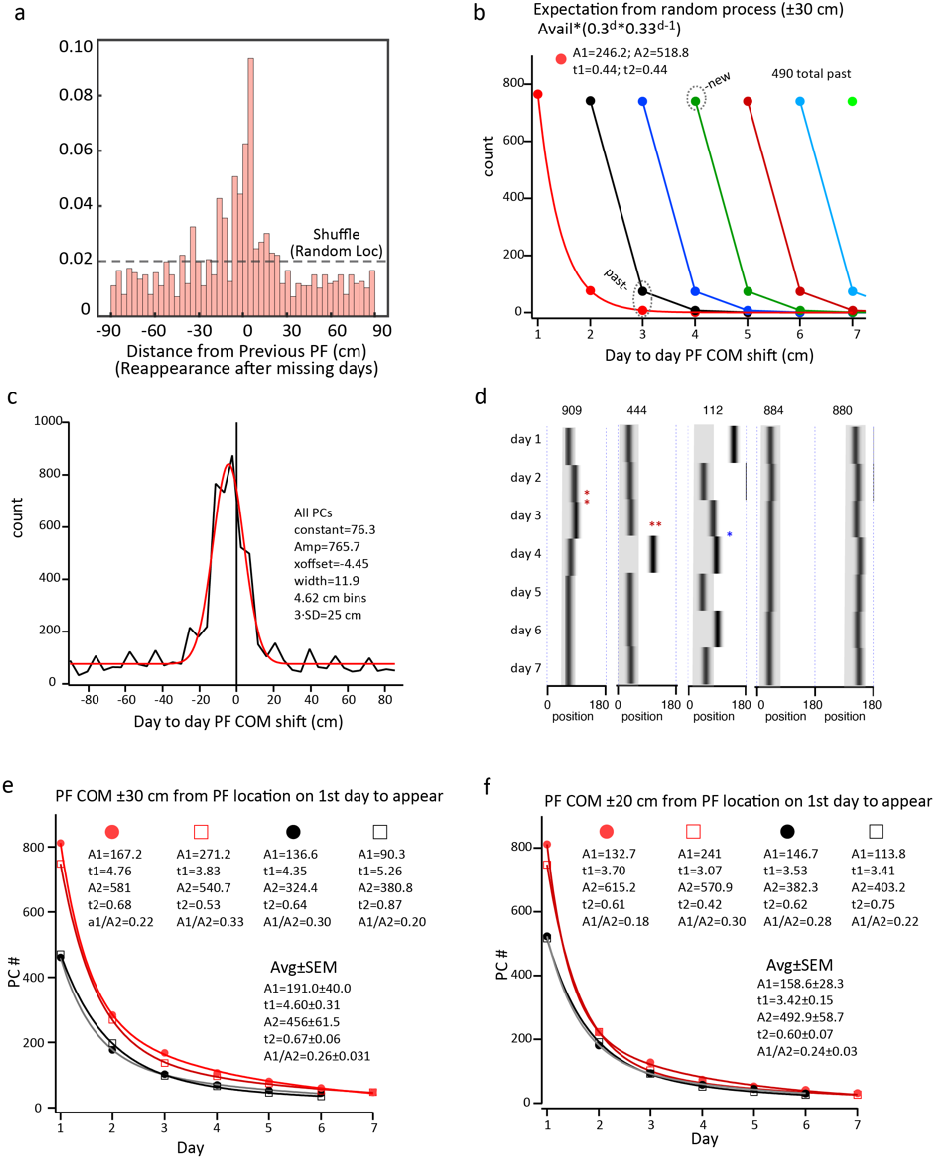
Probability of location for PC reappearance and determination of PF stability. **a**, Probability distribution for reappearance location of a previous PF after missing 1-3 days of activity as a function of the previous PF location (RL1 and RL2 data from 10 animals combined). Dashed line indicates expected distribution from random remapping after reappearance. **b**, Expected cell counts (plotted as in Fig. 1e) for a random process (i.e. no memory) where mean probability of any PC being active is 0.30 (mean PCs/total imaged pyramidal cells) and fraction of track for ±30 cm window (0.33). Count for given day = available neurons·(0.30^day^·0.33^day-1)^ where for cells first appearing on day 1 (red), d increments from 1-7. For cells first appearing on day 2 (black), d increments from 1-6, etc. Available neurons reduced by count of cells in past pool on each day. **c**, Distribution of day-to-day PF location shifts for all PCs from both reward location conditions (black). Gaussian fit to distribution (red) with listed parameters. **d**, PF locations for example PCs that had PF for 7 consecutive days in RL2. Cell 909 is example of PC that would have dropped out on day 3 (red asterisk) due to ±20 cm window (gray shade) but not ±30 cm window. Cell 444 is example of PC that would have dropped out on day 4 (two red asterisks) due to ±30 cm window (gray shade). Cell 112 falls outside of ±30 cm window every day except day 3 (blue asterisk). Cell 884 and 880 are inside of ±30 cm window all 7 days. **e**, Data from cells that appeared first on day1 and day 2 for each reward condition when using ±30 cm consistency criteria (as in fig. 1e). Parameters of double exponential fits are shown for each group. **f**, Data from cells that appeared first on day1 and day 2 for each reward condition when using ±20 cm consistency criteria. Parameters of double exponential fits are shown for each group.

**Extended Data Fig. 4:**
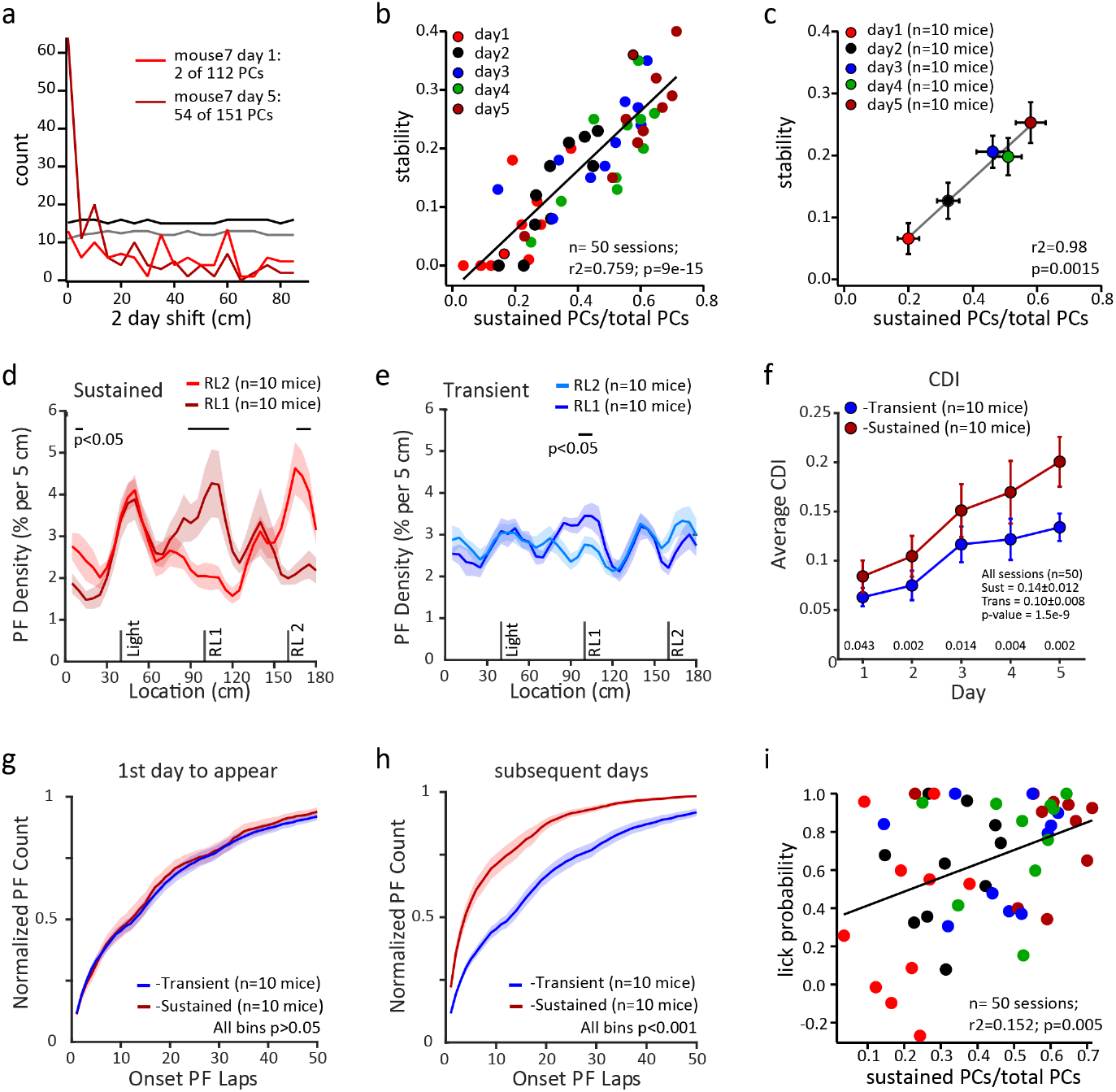
PC stability and animal-wise comparisons. **a**, Histogram of PF location shift from day 1 to day 3 (light red) with upper 99% CI (gray) and from day 5 to day 7 (dark red) with 99% CI (black) for mouse 7. **b**, Stability index for PC population versus sustained PCs fraction of total PCs for sessions on days 1-5 for 10 mice (total sessions n=50; 10 mice and 5 days). Means from sessions color coded by day. Line indicates linear regression with r-squared and p values listed. **c**, Mean stability index (n=10 mice) versus mean sustained PCs fraction of total for sessions on days 1-5 (error bars are SEM). **d**, PF distribution as percentage of sustained PCs for each RL condition, each bin is mean (±SEM) of 10 mice. **e**, same as d but for transient population. **f**, Cellular level discrimination index for days 1-5. Each point is mean CDI (±SEM) of 10 mice. P values from two-tailed, paired students t-tests are shown at bottom. **g**, Cumulative distribution of onset lap (first trial PC appeared) for the first days that PC appeared for sustained (days 1-5) and transient groups (days 1-6). Each bin is mean (±SEM) of 10 mice. **h**, Sustained group on subsequent days (2-7). Transient group is same as in panel (**g). i**, Lick probability versus sustained PC as fraction of total PCs for all 50 sessions. Each point is the mean from a single mouse color coded as above (**b**). Line indicates linear regression with r-squared and p values listed.

**Extended Data Fig. 5:**
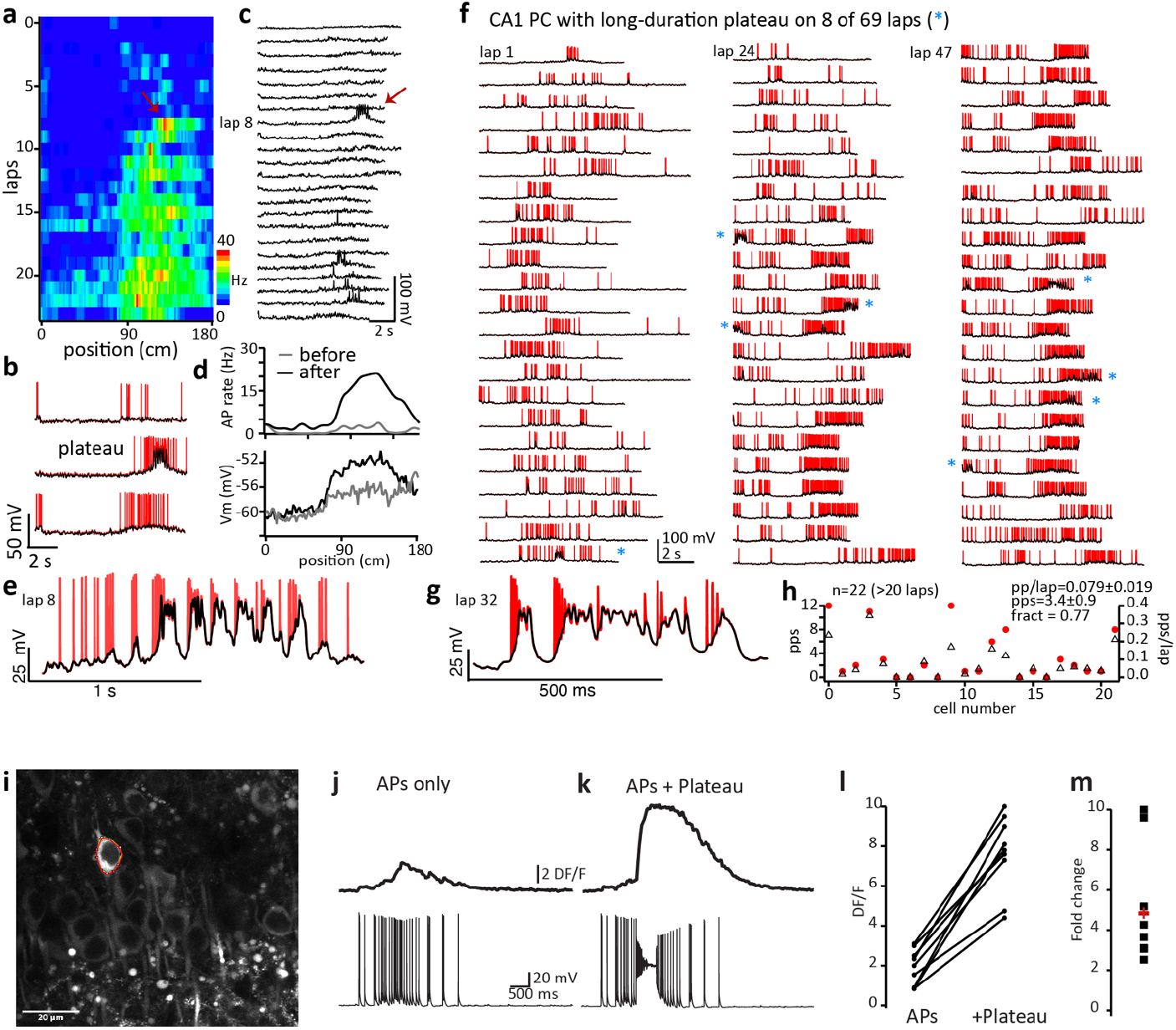
Dendritic plateau potential rates and GCaMP6 calibration. **a**, Firing rate heat map showing plateau potential initiation (red arrow) converts weak activity into strong PF (data from ref 32). **b**, Vm for consecutive laps (7, 8 & 9) where red are raw data and black is V_m_ trace with APs removed to show slow depolarization associated with plateau. **c**, AP removed V_m_ traces for all laps. **d**, Average AP rate and V_m_ versus position (100 bins) for before (gray) and after (black) plateau. **e**, expanded V_m_ from lap 8 showing long duration plateau. **f**, V_m_ from neuron after reward location switch showing the identification of 8 of 69 laps with long-duration plateau (single or cumulative duration during sequential theta cycles >150 ms). **g**, expanded V_m_ from lap 32 showing long duration plateau. **h**, population data from 22 CA1 neurons showing long-duration plateau potential counts (pps) and rate (pps/lap). The fraction of neurons showing at least one long-duration plateau (fract) is listed. **i**, *ex vivo* image of CA1 neurons expressing GCaMP6 (from GP5.17 mice). ROI from recorded neuron indicated (red circle). **j**, Ca^2+^ signal (upper) and Vm (lower) recorded from neuron during current injection train (2 nA for 2 ms) designed to reproduce activity of a large amplitude PF recorded in vivo^32^ (maximum interspike rate = 172 Hz; average rate = 37Hz for 500 ms period at peak). **k**, Ca^2+^ signal (upper) and Vm (lower) recorded from same neuron during current injection of AP train (2 nA for 2 ms) plus current step to evoke plateau (400 ms, 600 pA) designed to reproduce activity of a large amplitude PF with plateau potential. **l**, Population data of □-F/F for AP train w/o and with plateau. **m**, Fold change of Ca^2+^ signals between AP train without and with plateau.

**Extended Data Fig. 6:**
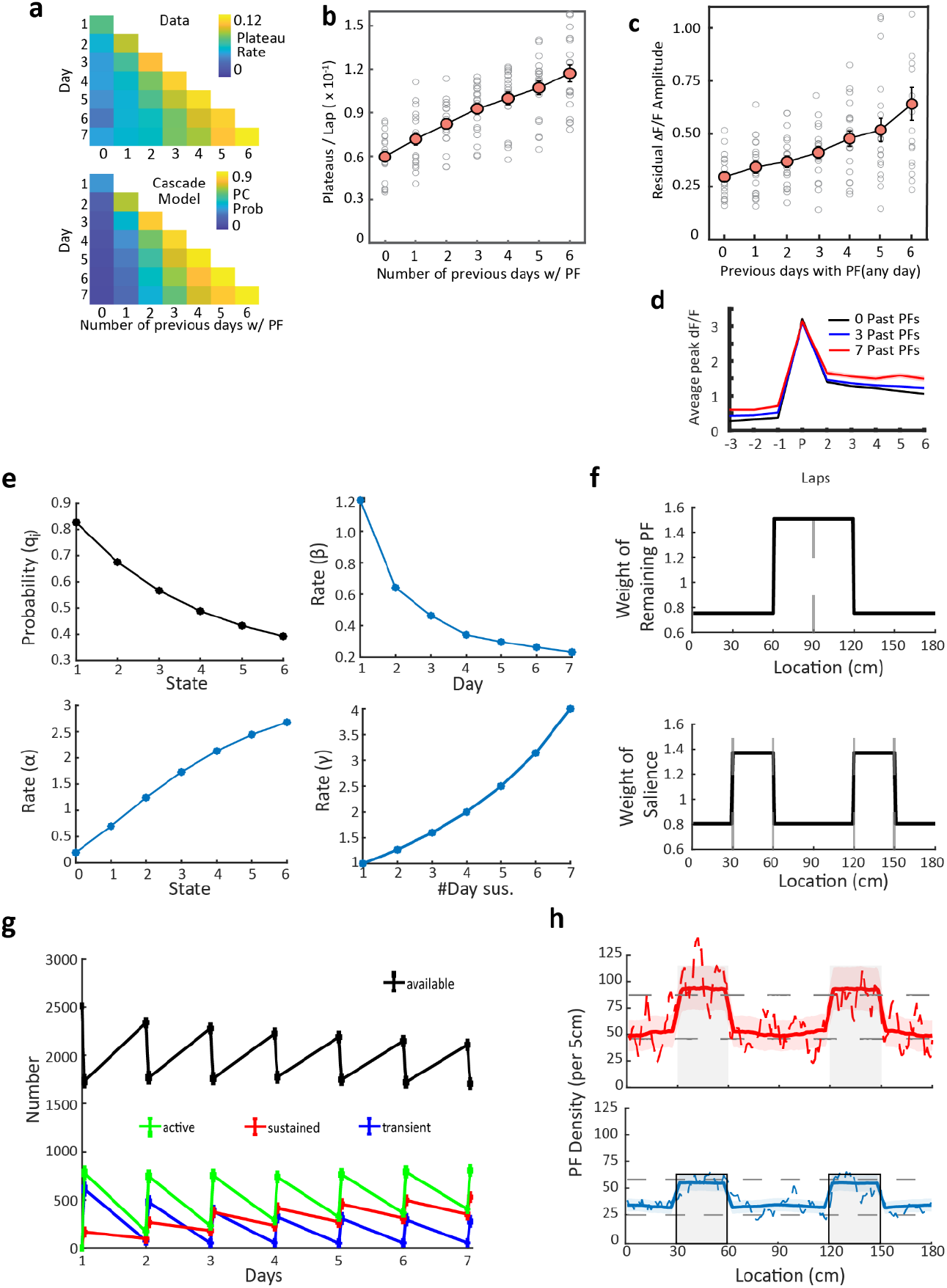
History dependent increase in putative plateau potentials, residual PC firing and parameters of cascade-type state model. **a**, Top, Average plateau rate per lap determined each day as a function of past PC activity from the experimental data. Bottom, same distribution determined as probability for PF formation from the red and blue pools in the cascade model (Fig.5c). **b**, Number of putative plateau potentials observed per lap sorted by the previous PF activity for the given PC (n=20, 10 animals, RL1 and RL2) (Combined data from all days). **c**, Amplitude of neuronal activity before BTSP induction event as a function of past PC activity in days. **d**, Abrupt establishment of a robust PC after a strong Ca^2+^ event showing increase in pre-event amplitudes with number of days of past PC activity (black, n= 1987 events; 0 past PCs) (blue, n= 1118 events; 3 past PCs) (red, n = 314 events; 6 past PCs). **e**, Visualization of the parameters of the cascade model. Top left: Probability of an active PF transitioning to the ‘unreliable’ resting state as a function of PF state (prior PF activity). Bottom left: Reactivation rate (α) of PFs as a function of PF state. Top right: Scaling factor for PF activation (β) as a function of PF state. Bottom right: Modulation factor (γ) for location specificity as a function of PF state. **f**, Illustration of factors influencing PF location specificity. Top: An example showing modulation by previously sustained PF with previous PF location at 90 cm and a modulation factor (γ) of 2. Bottom: An example showing modulation by salience with salience regions located between [30,60] cm and [120,150] cm. **g**, Count of model cells for transient (blue), sustained (red) and available (black) pools during a continuous model run for 7 days. **h**, PF densities for sustained PFs (top, red) and transient PFs (bottom, blue) estimated from 1000 model runs. Solid runs represent the average with the shaded area depicting the standard deviation. Dashed lines depict a single run example. Gray lines are 99% confidence intervals of random distribution produced by bootstrapping 10,000 time (averaged from 1000 runs). Sustained groups include identified PFs from days 1-7 and transient groups includes identified PFs from days 1-5.

**Extended Data Fig. 7:**
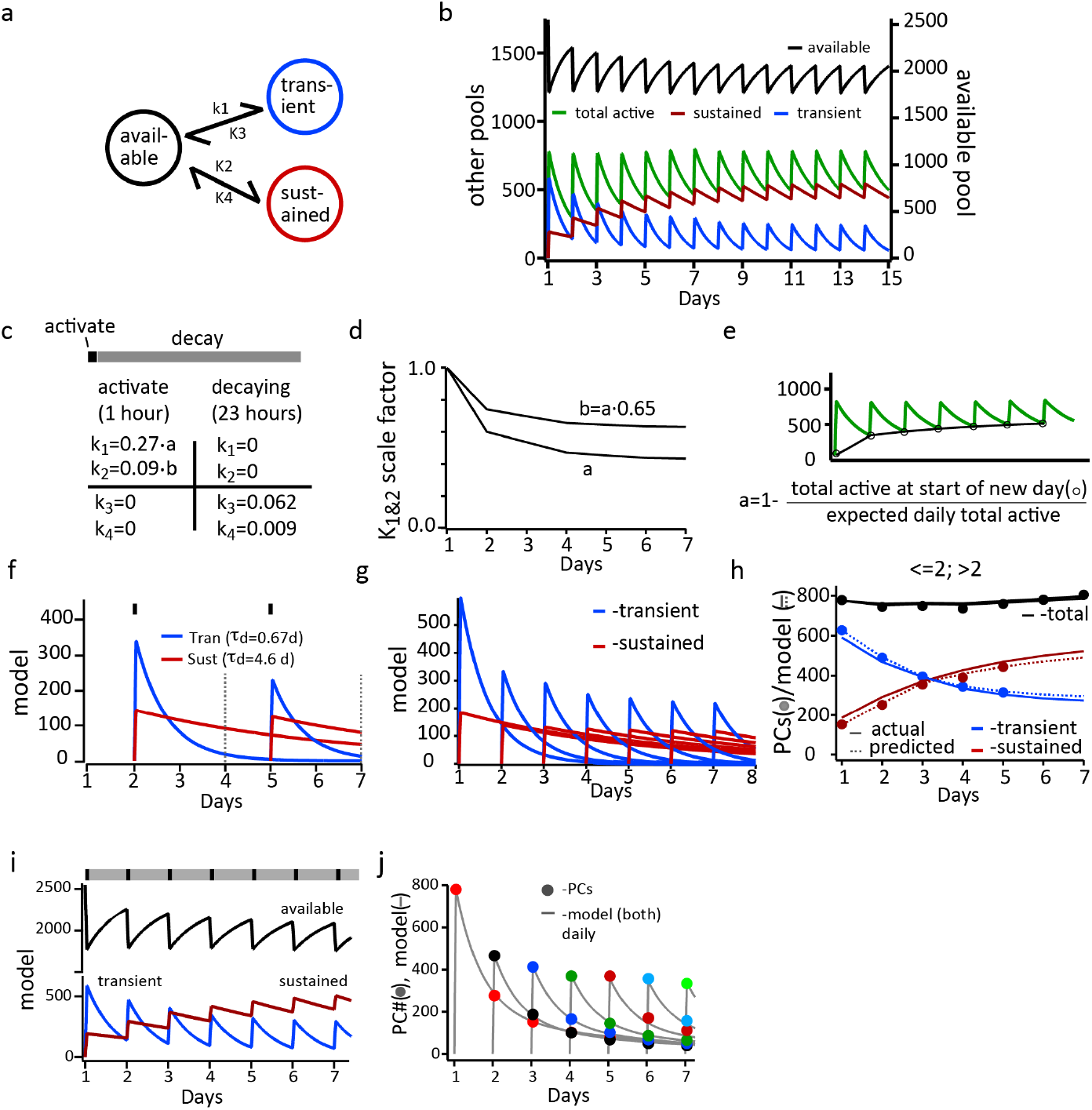
Three Pool Model. **a**, Schematic of model with three pools. **b**, Continuous model run for 14 days showing various pools. **c**, table showing model parameters used. **d**, K1, K2 scaling factors versus days. **e**, Calculation of scale factors. **f**, Single day runs based on continuous model showing transient (blue) and sustained (red) pools for days 2 and 5. Dashed lines indicate pool values on day where the actual data populations will be distinguished. **g**, transient and sustained components for each single day run. **h**, PC numbers for transient and sustained components from data (symbols). Solid lines are actual sustained and transient pools from model and dashed lines are estimates using the same procedure as used for data. **i**, Model cell counts for transient (blue), sustained (red) and available (black) pools for continuous model run for 7 days. At top, 1 hour on-track, (100 time-steps for activation; Black rectangles) and 23 hours off-track (2300 timesteps for decay; gray rectangles) are shown for each day. **j**, PC numbers (average of two RL conditions), color scheme same as panel Fig 2e. Gray lines are from single day runs of model with sustained and transient components combined.

**Extended Data Fig. 8:**
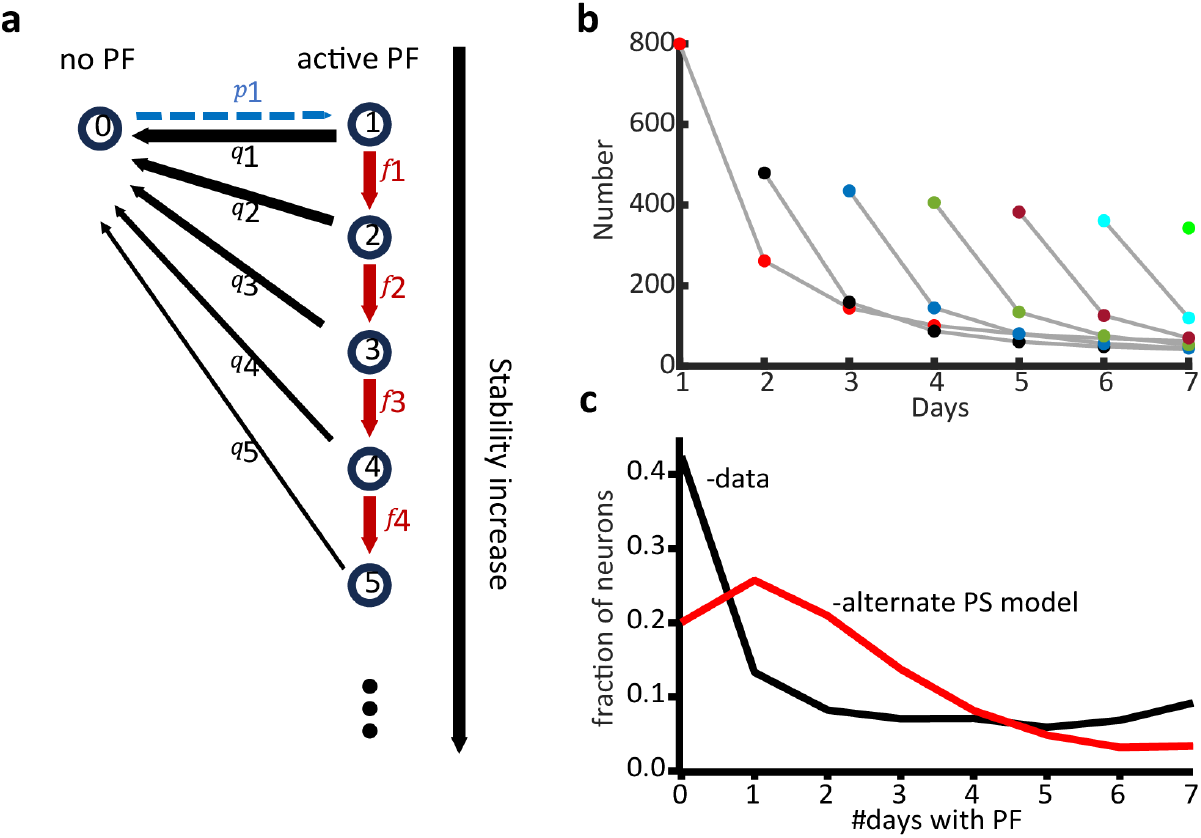
Alternative Progressive Stabilization Model. **a**, Alternate progressive stabilization (PS) model where decay rates (*q*) decrease with each day a PC is active also fits some aspects of data such as the double exponential decay of stable PC activity **b**, But fails to fit others such as **(c)** the number of days with a PF.

## Methods

### Mice and Surgery

All experiments were performed according to methods approved by the Baylor College of Medicine Institutional Animal Care and Use committee (Protocol AN-7734) and in compliance with the Guide for Animal Care and Use of Laboratory Animals. The data was collected from 29 GP5.17 mice of either gender (Jackson Laboratory, stock no. 025393). All experiments were performed in adult (>10 weeks) mice. Mice were housed under an inverse 12-hour dark/12-hour light cycle (lights off at 9 am) with temperature (∼21 degrees Celsius) and humidity (∼30-60%) control. All surgical procedures were performed under deep isoflurane anesthesia as described before^28^. After locally applying topical anesthetics, the scalp was removed, and the skull was cleaned. A 3 mm-diameter craniotomy was centered at 2.0 mm posterior from Bregma and 2.0 mm lateral from the midline above the hippocampus. Cortical tissue within the craniotomy was slowly aspirated under repeated irrigation with warmed sterile saline 0.9%. Once the external capsule was exposed, the cannula (3 mm diameter, 1.7 mm height) with a window (CS-3R, Warner Instruments) on the bottom was inserted and cemented to the skull. Finally, a custom-made titanium head bar was attached to the skull parallel to the plane of the imaging window using dental acrylic (Ortho-Jet, Lang Dental). Mice were given a recovery period of one week before any further behavioral training.

### Behavioral Apparatus

A linear treadmill apparatus with stationary head-fixation posts and a self-propelled belt of length 180 cm was used to train the animals to run and perform subsequent behavioral experiments. The treadmill was equipped with a rotary encoder coupled to an Arduino-based microcontroller to quantify the animal’s position and velocity. The location data was interfaced with a behavioral control system using a BPod module (Sanworks LLC) that delivered rewards through a solenoid valve (quiet operation, Lee valves) or light cues through an LED system at appropriate locations of the belt. All behavioral variables (position, velocity, lap markers, licks, trial types) were digitized at 10KHz via a PCIe-6343, X series DAQ system (National Instruments) using WaveSurfer software (wavesurfer.janelia.org).

### Behavioral Task and Training

After a 7-day recovery period, mice were placed on water restriction (1.5ml/day). The behavioral training started with habituation to the experimenter for water rewards for 30 mins per day for at least 5 days. The mice were then introduced to the treadmill and trained to run for water rewards at random locations on a blank belt with no sensory features. The mice were accustomed to the light cue (blue LED positioned in front of both eyes, flashing at 10 Hz for 500 ms) as well as 2-photon imaging during later parts of the training regime. The mice were trained for 5-7 days and had to run 100 laps in one hour before they were deemed ready to be introduced to the experimental task.

On day 1 of the behavioral task, the mice were introduced to a new featured belt (6 varying sensory cue patterns of approximately 15 cm equidistantly placed on a new belt). A light cue on either side of the animal was triggered at 40 cm (flashing 10 Hz for 500 ms) that predicted the reward location at 100 cm when activated on the left side and 160 cm when activated on the right side. Trials for each location were grouped in epochs of 12-18 laps and randomly switched. Mice performed one session per day of this task for a succession of 7 days.

### Two-photon Ca^2+^ longitudinal image acquisition and signal processing

All *in-vivo Ca*^*2*+^ imaging recordings were performed in the dark using a custom-made two-photon microscope (Janelia MIMMS design). Transgenetically expressed GCaMP6f was excited at 920 nm (typically 40 – 60 mW) by a Ti:Sapphire laser (Chameleon Ultra II, Coherent) and imaged through a Nikon 16x, 0.8 Numerical Aperture (NA) objective. Emission light passed through a 565 DCXR dichroic filter (Chroma) and was detected by GaAsP photomultiplier tubes (11706P-40SEL, Hamamatsu). Images (512 × 512 pixels) were acquired at ∼30 Hz using the Scan-Image software (Vidrio Technologies, LLC).

A reference field of view (FOV) was chosen and registered before the start of the experiment for every animal. Each day, the FOV was aligned to this reference and the experiment aborted if differences were noted in the imaging plane on subsequent days. Acquired two-photon images were motion-corrected using Suite2p^59^ (Python version, https://github.com/MouseLand/suite2p). Images were registered across days and only data from mice with stable FOVs across 7 days was considered for further processing. Image Segmentation into regions of interest (ROIs) to identify neuronal somata was done manually in ImageJ (version 2.3.0) to ensure that each ROI could be reliably tracked across all days. Briefly, frames were subsampled from across all sessions to form a composite image across days. Image segmentation was performed on this composite image with subsequent manual verification that every ROI adequately and exclusively represented the given cell on each day. Signal extraction was performed using custom code in Python (scipy.ndimage).

The raw fluorescence was converted to Δf/f, where Δf/f was calculated as (F – F0)/F0, where F0 is the 50^th^ percentile of a 25-s moving window. Only significant Ca^2+^ transients, defined as transients larger than 3 Standard Deviations above the baseline noise were considered as functional activity for any further analysis. Baseline noise was estimated from deviations below peak histogram values of all Δf/f activity.

### Determination of Place Field Activity

Spatial maps of neural activity were formed by dividing the 180 cm track into 50 spatial bins of 3.6 cm each. For each spatial bin, for every lap, the mean Δf/f was calculated when the velocity of the animal was above 2cm/s.

A behavioral epoch was determined to have place field (PF) activity if 1) Spatial Activity in the epoch had spatial information that exceeded 95% confidence interval determined by shuffling the epoch activity as previously described^60^. 2) Only laps with peak Δf/f within 54 cm (15/50 bins) of the peak average epoch activity were considered for further PF analysis. 3) An onset lap could be determined as the first instance where 3/5 laps showed PF activity. 4) The reliability of PF activity after the onset lap was 40% for the rest of the epoch.

The PF location for a given reward location was determined as the peak of the average lap activity for all laps within the active behavioral epochs for that reward location.

### Determination of day-to-day Place Field stability criteria and stability index

To determine the amount of day-to-day PF shifting or jitter that should be tolerated for a PC to be considered as having a consistent PF we generated a distribution of the day-to-day PF shifts of the population of PCs. This distribution was generally normal (however, with a slight negative shift from zero) and thus was fit with a gaussian function from which 3 SD was found to be 25 cm (Extended Data Fig. 2a). Based on this we used a window of ±30 cm to fully capture the PCs within this central process. Thus, to be considered a stable PC the neuron had to express a consistently active PF on each consecutive day as defined by its PF location being within 30 cm of that on the first day of appearance. We calculated the expected results (as in Fig. 2a) of a random, memory-less, process that used the ±30 cm window of PF jitter (A·0.3^d^·0.33^d-1^). Where A is the available population of neurons on a given day (starts at 2511 on day 1 and decreased as A minus pastPCs on a given day), 0.3 is the probability of any particular neuron becoming a PC (given the fraction of average PC count and total imaged neurons or 767/2511), 0.33 is the jitter window (60 cm/180cm) and d is the day of the “recording”. The difference between the analysis shown in Extended Data Fig. 3c and that in Fig. 2e gives confidence that using the ±30cm window does not cause our data to be heavily impacted by a random process. Finally, we analyzed the PC data using a ±20 cm window and found that although there were fewer PCs in the stable populations the data were still well fit by a double exponential with similar time constants as found using the ±30 cm window (Extended Data Fig. 3e,f). We surmise that the tighter window slightly reduced the time constants as more PCs were removed from the stable groups on later days because they had shifted outside the window for one day during the week (even if they returned the next day; see Extended Data Fig. 3c, cell 909). In this way the tighter time window can be viewed as overly restrictive.

For calculations of sustained and transient PC numbers (Fig. 2) we identified transient cells as those that had a stable PF for 1 or 2 days (≤2 days) and sustained as those with a stable PF for 3 or more days (>2 days). For calculations of sustained and transient PC properties (Figs. 3 &4) we identified transient cells as those that had a stable PF for 1 day (1 day) and sustained for 3 or more days (>2 days).

PC stability index was the fractional difference between the number of PCs per mouse that had a minimum level of shift (<20 cm) and that expected from random. To calculate this we subtracted the upper 99% confidence interval of a random distribution from the total number of PCs with a two-day interval (d versus d+2) shift <20 cm (1^st^ three 5 cm bins) then divided this by the total number of PCs in that mouse for the given day (Extended Data Fig. 4a).

### Behavioral data quantification

The velocity and licking behavior was mapped into 50 spatial bins per running lap for quantification of behavior. For licking selectivity, the probability of licking at a given location bin was quantified as the percentage of laps with at least one lick inside the given spatial bin for the entire session. The reward anticipatory zone was defined as 3 bins before reward location and the random zone consisted of 3 bins not overlapping the active or alternate reward zone around the light stimulus. Licking Selectivity was quantified as

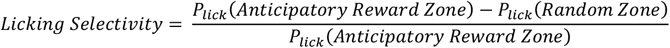

### Trial-by-trial Cosine Similarity and Discrimination Indices

Cosine Similarity Matrices for individual cells and across population were calculated as previously described^13^. Briefly, for the Cosine Similarity Matrix of an individual cell, every row of the cell’s spatial activity map, *Ã*, was divided by its *l*_2_ norm to give the matrix 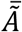. The cosine similarity matrix for each cell is then given by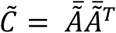. For Population Similarity matrices, the single cell spatial activity maps were horizontally concatenated to form a fat matrix *A* = [*A*_1_|*A*_2_| … |*A*_*N*_], where *N* is the number of neurons recorded in that session. Each row of *A* was divided by its *l*_2_ norm to give the matrix Ā. The Population Similarity Matrix was then given by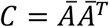.

The Cellular Discrimination Index was calculated for each cell from 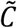. For each row of 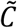, the average ‘within RL’ cosine similarity per row was calculated as the average value from all trials of the same reward location and ‘between Rl’ cosine similarity per row, as the average value from all trials of the alternate reward location. The difference in the within RL and between RL averages across all rows was deduced as the Cellular Discrimination Index (CDI). The Population Discrimination Index (PDI) was likewise calculated using the Population Similarity Matrix, *C* as the starting matrix.

### BTSP and Plateau Analysis

A putative BTSP event was identified with the following criteria: 1) It was a strong Calcium event with amplitude in the top 20^th^ percentile for all Calcium events in that cell for a given session with a minimum amplitude of 1 Δf/f. 2) Only laps with peak Δf/f within 45 cm of the putative BTSP event peak were considered for further BTSP analysis. 3) The event was associated with 3/5 subsequent laps actively firing, and 4) the average COM of the post-firing laps had a negative shift compared to the putative BTSP event. This first events in a session for each context identified with these simple criteria showed other independent signs of BTSP viz, a) an abrupt increase before and after, in the PF amplitude and b) a strong correlation of induction velocity during the event and PF width (Fig.4).

The Calcium signal threshold for a plateau potential was the minimum of all identified BTSP events (if any) detected during the given session for that context in a given cell. We provide independent measurements of Δf/f amplitude for PF-like firing with and without identified Calcium plateau events in ex vivo physiology experiments and that of in vivo plateau rate in awake behaving mice using whole cell patch clamp recordings to corroborate the Δf/f plateau threshold values and plateau rates observed during in vivo Calcium imaging using the above criteria (Extended data fig. 5).

### In Vitro whole-cell methods

400 µm transverse hippocampal slices were cut from 8-16 week old male and female GP5.17 GCaMP mice using Leica Vibratome VT1200S after perfusing an isoflurane-anesthetized animal with ice-cold solution containing in mM: 210 Sucrose, 25 NaHCO_3_, 2.5 KCl, 1.25 NaH_2_PO_4_, 0.75 CaCl_2_, 7 MgCl_2_, 7 Glucose. Slices were incubated for 30 mins at 35ºC in, and then recorded from at 35ºC, in a solution containing: 125 NaCl, 25 NaHC0_3_, 3 KCl, 1.25 NaH_2_PO_4_, 1 MgCl_2_, 2 CaCl_2_, and 16 dextrose. All solutions contained fresh Na-pyruvate (3 mM) and ascorbic acid (1 mM), and were bubbled with 95% O_2_ and 5% CO_2_. To enhance plateau initiation 250 nM muscarine was bath applied after recording began. Cells were visualized using an Olympus BX-61 microscope using a water-immersion lens (60X, 0.9 NA, Olympus, Melville, NY). Whole cell current clamp recordings were performed using a Dagan BVC-700 (Minneapolis, MN) in active “bridge” mode and analog-filtered at 10 kHz before being digitized at 40kHz. The pipette solution was the same as described for the in-vivo electrophysiology but also included 20–40 µM Alexa 594 (Invitrogen). Attempts were made to maintain R_series_ above 20 MΩ to avoid excessive dialysis. Imaging was performed using a resonant galvanometer-based two-photon laser scanning system (Ultima; Bruker Technologies, Middleton, WI; using Chameleon Ultra II; Coherent, Auburn, CA). Full field scans for Ca^2+^ imaging were performed at 30 Hz using excitation at 920 nm. Repetitive APs without plateau were elicited by a train of current injections (2 nA, 2 ms) that mimicked a strong PF recorded in vivo^34^. Plateaus were added by including an additional 500 ms, 400-600 pA current step in the middle of the AP train.

### In Vivo whole-cell methods

The intracellular recordings were performed as previously reported^32-34,61^. Data collected for Qian et al (ref 54) and reanalyzed here. Briefly, an extracellular LFP electrode was firstly lowered into the dorsal hippocampus using a micromanipulator until prominent theta-modulated spiking and increased ripple amplitude were detected after passing through neocortex, usually to a depth of 1.0–1.2 mm. Then a glass intracellular recording pipette was lowered to the same depth while applying positive pressure (∼9.5 psi). The intracellular solution contained (in mM): 134 K-Gluconate, 6 KCl, 10 HEPES, 4 NaCl, 0.3 MgGTP, 4 MgATP, 14 Tris-phosphocreatine, and 0.2 % biocytin. Current-clamp recordings of intracellular membrane potential (V_m_) were amplified and digitized at 20 kHz, without correction for liquid junction potential.

Mouse behavior was as above except that reward switch occurred once and only tactile cues were present. To analyze the spatial location of AP rate, the spatially binned AP rate was determined (AP#/time in bin) using the spatial bin size of 1.8 cm for each trial and averaged over the duration of the recording. To analyze V_m_ ramps, APs were removed by deleting all points 0.26 ms before and 3 ms after a threshold value (dVm/dt = 25 V/s), resulting traces were linearly interpolated, baseline corrected on a trial-by-trial basis by subtracting the difference in the recorded AP threshold (the most negative 5^th^ percentile) and –50 mV and spatially binned and averaged as above. The spatially-binned AP rates and V_m_ ramps in the heatmap were smoothed with a Gaussian of 5 bins (5 cm) and 11 bins (11 cm), respectively. Spontaneous naturally-occurring plateau events were detected from AP-removed traces as events where V_m_ crossed -35 mV and duration determined as the time above this threshold. These events were counted as long-lasting plateau if the duration of a single plateau potential was longer than 150 ms or if the duration of multiple plateaus occurring on consecutive theta cycles (threshold crossings were separated by <150 ms) was longer than 150 ms (Extended Data Fig.5).

### 3-pool Computational model

To simulate the production of two groups of active PCs with distinct PF activity decay rates we used a standard first order competitive interaction between three cellular pools, which modelled (1) available pyramidal cells, (2) transient cells and (3) sustained cells, as schematized (Extended Data Fig. 7). The reactions were separated into two phases to reproduce the two components of the behavior (time on- and off-track). Forward rate equations (activations) were:

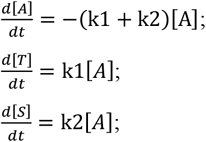

where initial [A] was 2511 and dt was 0.01 hour for 100 time steps (1 hour). After which the elements were allowed to decay back to the available population for an additional 2300 time steps (23 hours) according to rate equations (decay)

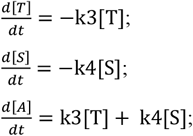

Rate constants (k1, k2, k3, k4) are shown in Extended Data Fig. 7. The values of each of these were determined from the data where k3 and k4 were set by the fast and slow time-constants, respectively, produced in Fig. 2. [A] was initialized to the total number of imaged neurons. k1 and k2 by the initial, fractional amplitude of the slow component (Amp2) determined in Figure 1. Initial [A] = 2511

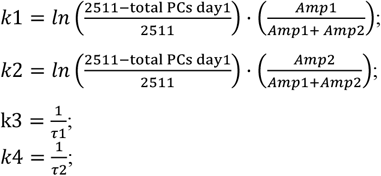

To reproduce multiple days of on- and off-track experience, repeated activations and decays were given and k1 and k2 were scaled daily by the fraction of the population present at the beginning of each day and the desired daily total population from data

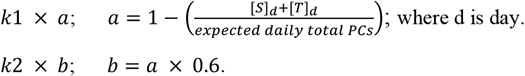

b was the only free parameter in the model and was adjusted to minimize MSE between the model results and the data shown in Fig. 2e.

To generate the data to compare with that in Fig. 2e we ran the model for single day activations, starting at day 1 and progressing through day 7. The daily values of k1, k2 and [A]t calculated from the multiple day run of the model were used. To determine how well the model corresponded to the data we calculated residuals (total residual= -10.1; n=28) and mean squared error (MSE= 182; n=28) between the model results and the 28 data points shown in Fig. 2e (see Extended Data Fig. 7j).

### Cascade-type state model

The cascade model (Figure 5c) classifies the cells in a series of states determined by the number of days a cell has exhibited an active place field. For instance, a state denoted as ‘3’ implies that the cell has shown an active place field for three days, which need not be consecutive or at the same location. It further divides the cells in resting state (between sessions) into two distinct groups: those that reliably reappear at the same location in the next session, ‘reliable PFs’(red), and those that probabilistically reappear, ‘unreliable PFs’ (blue). The transitions between different states within the cascade model are conceptualized as discrete, one-step jumps occurring on a daily basis. Similar to the three-pool model, each day is composed of two distinct phases: the ‘activation’ phase and the ‘decay’ phase. At the start of the simulation, the cells are initialized 88% to unreliable state ‘0’ and the remaining 12% to unreliable state ‘1’ to mimic the potential training effect on memory formation. During the ‘activation’ phase, unreliable cells at a given state ‘i’ can transit to a state ‘i+1’ as an active PF with probability *p*_*i*_. This transition, represented by a blue arrow in the diagram, involves the acquisition of a random place field location, influenced by the salience and historical context of place fields (see below). Concurrently, reliable cells at state ‘i’ progress to state ‘i+1’ as an active PF while maintaining their previous place field with a probability *f*_*i*_ (assumed to be 1).

In the decay phase, active PFs at state ‘i’ transition with a probability *q*_*i*_ to unreliable PFs at state ‘i’, while the remaining proportion (1 − *q*_*i*_) transit to reliable PFs at state ‘i’ (black arrows). Notably, *q*_*i*_ decreases as the state ‘i’ increases (Extended Data Fig.6e), reminiscent of previous work on cascade models of synaptic memory.

This *p*_*i*_ pathway plays a pivotal role in modulating cell representations based on salience and the history of PF. For any given day k, unreliable PFs at state ‘i’ have a probability *p*_*i*_ to transition into a place cell. The formula of *p*_*i*_ is given as

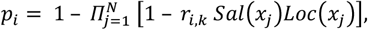

where,

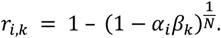

Here N is the number of spatial bins and *x*_*j*_ = *j* ∗ *L/N* denotes the location of the jth spatial bin. It is noteworthy that when both *Sal*(*x*_*j*_) and *Loc*(*x*_*j*_) equal to 1, *p*_*i*_ then equals to *α*_*i*_*β*_*k*_.

The PF location of these appearing place cells is determined randomly from N spatial bins,

following a specific probability distribution described by the formula:

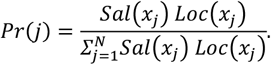

The rate *α*_*i*_ represents the likelihood of a plasticity event occurring for a cell at state ‘i’. This probability rate increases with the state (Extended Data Fig.6e). It implies that cells with a stronger history of active PFs in previous days have a higher probability of becoming a PC on the current day. Furthermore, the rate *β*_*k*_ serves as a global modulation factor that influences the frequency of plasticity events on a day-to-day basis which maintains a nearly constant number of place cells every day. This factor is analogous to the factors ‘a’ and ‘b’ in the three-pool model and is shown to decrease over time as more sustained place cells are established, as depicted in (Extended Data Fig.6e).

The probability of a plasticity event and its dependence on the animal’s location are further modulated by salience, following the formula:

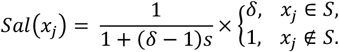

where *S* is the range of salience region, *s* is the ratio of the salience region compared to the total lap length, and *δ* denotes the modulation factor by the contribution of salience. δ is chosen to be 1.7 in this simulation. An illustrative example of this modulation by salience in regions between [30,60] cm and [120,150] cm is presented in Extended Data Fig.6f. In addition, the history of previously sustained PF also influences this probability, governed by the formula:

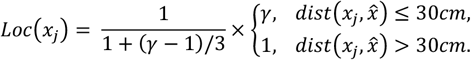

where 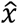 is the location of the previously sustained PF. An example showcasing the modulation effect for a previous PF with location at 90cm and *γ* equals to 2 is displayed in Extended Data Fig.6f. Notably, the amplitude of modulation *γ* varies depending on the duration for which the previous PF was sustained (within a limit of a 30 cm shift per day). The relationship between the degree of PF sustained-ness and modulation amplitude is detailed in Extended Data Fig.6e.

The model parameters are set up with the following formulas:

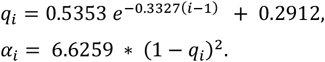

These parameters were obtained through the following steps: First, we initially estimated the transition probabilities of cells between active and non-active states, as well as the probability of maintaining the same place field location across different days. These estimations were conditioned on the histories of place field activity and locations derived from experimental data. Second, we fit the model parameters to best reproduce these transition probabilities. The value of location modulation parameter, *γ*_*l*_, is set up with the following formula:

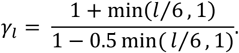

This formula ensures that the *Loc*(*x*_*j*)_ within the previous field 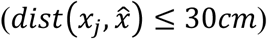increases as a threshold linear function of *l*, the number of days the PC stays at the same location. The value of daily modulation parameter, *β*_*k*_, is set slightly different for each run of the simulation and keeps the expected number of place cells every day consistent with the experiment data. The average value of *β*_*k*_ satisfies the formula:

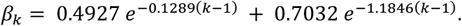

### Alternate progressive stabilization (PS) model

Another multi-pool model where the stability of place fields increases with consecutive days of presence is also studied (Extended Data Fig.8). We categorize cells into two groups: those without a detectable place field (PF) within a given day, termed “no PF”, and those exhibiting a detectable PF, referred to as “active PF”. For “active PF”. Cells are further classified into different states based on their stability, which are quantified by the number of days a cell has exhibited an active and consistent place field. For instance, a cell in state “3” has maintained an active place field at the same location for three days.

The transitions between different states within the model are also conceptualized as discrete, one-step jumps occurring daily, similar to the cascade model. The transitions are illustrated through directed arrows in the model diagram in Extended Data Fig.8a. Each day is composed of two distinct phases: the “activation” phase and the “decay” phase. During the “activation” phase, cells lacking a place field (“no PF”) possess a probability *p*_1_ to transit to the state “1” as a place cell (“active PF”). This transition, represented by a blue arrow in the diagram, involves the acquisition of a random place field location. Simultaneously, cells already exhibiting a place field (“active PF”) at state “i” can progress to state “i+1” while maintaining their place field with a probability *f*_*i*_ (assumed to be 1), as depicted by the red arrows. The *p*_1_ is set to be 0.3186 × β_*k*_ (where *k* is the day index) with the rate β_*k*_ serving as a global modulation factor to ensure a constant number of place cells across days.

In the decay phase, cells showing a place field within the session (“active PF”) at state “i” have a probability *q*_*i*_ (illustrated by a black arrow) to revert to the state of “no PF” without a detectable place field. Notably, *q*_*i*_ decreases as the state “i” increases as a power function: *q*_*i*_ = 0.67^*i*^, which aligns with the observed number of PCs lost after one session. This formula suggests that the stability of place cells increases as they persist in their activity across days, reminiscent of previous works on cascade models of synaptic memory^36,38,39^.

### Statistical methods

The exact sample size (n) for each experimental group is indicated in the figure legend or in the main text. No statistical methods were used to predetermine sample sizes, but our sample sizes are similar to those reported in previous publications^25,29,62^ using a similar behavioral task and by the expected number of active neurons that can be imaged with the two-photon microscope in awake behaving mice. Normality for data was tested with a Kolmogorov-Smirnov test before using any parametric statistical testing. If not otherwise indicated in the figure, data are shown as mean ± SEM.

